# Interactions established by isoform-specific TrkB-T1 sequences govern inflammatory response and neurotoxicity in stroke

**DOI:** 10.1101/2024.11.13.623346

**Authors:** Lola Ugalde-Triviño, Gonzalo S. Tejeda, Gema M. Esteban-Ortega, Margarita Díaz-Guerra

## Abstract

Ugalde-Triviño et al. develop cell-penetrating peptides derived from neurotrophin receptor TrkB-T1 to identify isoform-specific protein interactions and demonstrate protective effects on neuroinflammation and neurotoxicity reducing brain damage in a mice model of ischemic stroke, of relevance to human therapy.

**Abstract:** Glia reactivity, neuroinflammation and excitotoxic neuronal death are central processes to ischemic stroke and neurodegenerative diseases, altogether a leading cause of death, disability, and dementia. Due to the high incidence of these pathologies and the lack of efficient treatments, it is a priority developing brain protective therapies impacting both neurons and glial cells. Truncated neurotrophin receptor TrkB-T1, a protein produced by all these cells, plays relevant roles in excitotoxicity and ischemia. We have hypothesized that interactions established by isoform-specific TrkB-T1 sequences might be relevant to neurotoxicity and/or reactive gliosis and, therefore, constitute a therapeutic target. We identify here the TrkB-T1-specific interactome, poorly described to date, and demonstrate that interference of these protein-protein interactions using brain-accessible TrkB-T1-derived peptides can prevent reactive gliosis and decrease excitotoxicity-induced damage in cellular and mouse models of stroke. The pivotal role played by TrkB-T1 on microglia and astrocyte reactivity suggests that isoform-derived peptides could become important in development of therapies for human stroke and other excitotoxicity-associated pathologies.

## Introduction

Stroke is a leading cause of death, disability, and dementia. The ischemic type (≈85% of total cases) is produced by blockage of a brain blood vessel and decreased levels of nutrients and oxygen in the affected region, which result in neuronal death and irreversible brain damage. After an insult, two damaged areas can be distinguished, the infarct core and the penumbra. The first one contains irreversible damaged tissue while the penumbra is functionally impaired but metabolically active and, therefore, could be recovered. Accordingly, the penumbra has become a suitable and interesting target for neuroprotection, although, if no therapy is applied, neurons in this area can suffer secondary neuronal death. This process is mainly due to excitotoxicity induced by overactivation of glutamate receptors, primarily those of the N-methyl-D-aspartate (NMDARs) type, followed by massive Ca^2+^ entry and loss of ion homeostasis. Another important contribution to ischemic stroke pathophysiology is the inflammatory response (DeLong et al., 2022). In the acute phase, resident immune cells including astrocytes and microglia are activated, followed by passage of circulating immune cells across damaged blood–brain barrier (BBB) to reach the injured tissue. Reactive astrogliosis, a process taking place in the peri-infarct environment, implies a dramatic astrocyte transformation that increases proliferation and migration, modifies morphology and leads to upregulation of many genes, including that encoding glial fibrillary acidic protein (GFAP) (Shen et al., 2021). A subtype of reactive astrocytes loses homeostatic properties and experiences a pro-inflammatory gain of function with detrimental effects on neuronal and oligodendrocyte survival (Escartin et al., 2021; Hasel et al., 2021; Verkhratsky et al., 2021). Neuronal function is also regulated by microglia through release of neurotrophic factors or by specialized junctions formed between microglia processes and synaptic elements or neuronal cell bodies (Cserep et al., 2020). Interestingly, a cross-talk between astrocyte and microglia activation is established in the central nervous system (CNS) and, for example, neurotoxic reactive astrocytes are induced by microglia secretion of interleukin 1α (IL-1α), tumor necrosis factor (TNF), and complement component 1, subcomponent q (C1q) (Liddelow et al., 2017).

The molecular mechanisms underlying excitotoxicity and reactive gliosis are still largely undefined even though a better knowledge of them might help to identify new targets for stroke therapy. Nowadays, conventional pharmacological and mechanical treatments directed to recover tissue reperfusion have relevant limitations and reach a minority of patients (reviewed by Prabhakaran et al., 2015). Therefore, we need to identify new candidates for stroke therapy and develop strategies that ideally target the different pathological processes and cell types involved. A signaling pathway that becomes profoundly aberrant in ischemic stroke is mediated by binding of brain-derived neurotrophin factor (BDNF) to its high-affinity transmembrane receptor, tropomyosin-related kinase B (TrkB) (Vidaurre et al., 2012; Tejeda and Diaz-Guerra, 2017). In physiological conditions, BDNF is produced by multiple cells, including neurons and glia (Riley et al., 2004), and promotes neuronal maturation, differentiation, survival and synaptic function, among other processes. Different isoforms of the TrkB receptor are produced by alternative splicing of the *Ntrk2* gene (Lei and Parada, 2007), full-length TrkB (TrkB-FL; 145 kDa) and truncated TrkB-T1 (95 kDa) being the major isoforms in the murine cortex (Middlemas et al., 1991). TrkB-T1 is the predominant isoform produced in the adult mammalian nervous system where it is mostly expressed by astrocytes and, secondarily, neurons. In contrast, TrkB-FL is almost exclusively expressed in neurons (Niu et al., 2024). There, BDNF binding to TrkB-FL induces receptor dimerization, increased tyrosine kinase (TK) activity and receptor transphosphorylation, leading to activation of interconnected signaling pathways that, altogether, stimulate different processes central to CNS functioning. They include activation of the prosurvival transcription factors (TFs) cAMP response element-binding protein (CREB) (Bonni et al., 1999) and myocyte enhancer factor 2 (MEF2) (Liu et al., 2003; Wang et al., 2007), which regulate a wide spectrum of genes including those encoding BDNF (Lyons et al., 2012; Shieh et al., 1998; Tao et al., 1998), TrkB (Deogracias et al., 2004) or certain NMDAR subunits (Lau et al., 2004). In neurons, BDNF can also bind to truncated TrkB-T1, isoform lacking the TK region and traditionally considered as a dominant-negative receptor that inhibits BDNF-signaling by forming heterodimers with TrkB-FL (Biffo et al., 1995). TrkB-T1 expressed in hippocampal astrocytes also participates in fine-tuning of neurotrophin signaling by BDNF storage and translocation after internalization of BDNF/TrkB-T1 complexes (Alderson et al., 2000).

Receptor TrkB-T1 also regulates important astrocyte functions in response to BDNF, including calcium release from intracellular stores (Rose et al., 2003), transport of glycine (Aroeira et al., 2015) and gamma aminobutyric acid (GABA) (Vaz et al., 2011), and cell maturation (Holt et al., 2019) and morphology (Ohira et al., 2005). It has been proposed that these TrkB-FL-independent actions rely on TrkB-T1 interactions established by a highly conserved and isoform-specific C-ter sequence (FVLFHKIPLDG) (Biffo et al., 1995; Middlemas et al., 1991). Interestingly, TrkB-T1 is also produced outside the nervous system (e.g. heart, pancreas) where it plays important physiological actions (Fulgenzi et al., 2020; Fulgenzi et al., 2015). Anyway, the only TrkB-T1-interacting protein identified so far is Rho GDP dissociation inhibitor 1 (RhoGDI1), found in astrocytes (Ohira et al., 2005). By sequestering Rho GTPases in a GDP-bound state in the cytosol, RhoGDI1 controls this important family of regulatory proteins that, among others, link surface receptors to cytoskeleton organization, regulation of gene transcription and neuronal survival/death (Stankiewicz and Linseman, 2014). Binding of BDNF to TrkB-T1 in astrocytes disrupts constitutive TrkB-T1/RhoGDI1 complexes and releases Rho-GDI1 protein which then regulates Rho GTPases (Ohira et al., 2005).

The expression of TrkB-T1 is upregulated in multiple neurological disorders, including stroke, spinal cord injury, several neurodegenerative diseases (NDDs) or Down syndrome (reviewed by Cao et al., 2020). In experimental stroke, TrkB-T1 levels increase in astrocytes surrounding the infarct core while neuronal TrkB-FL decreases inside this region (Ferrer et al., 2001). Additionally, astrocyte TrkB-T1 signaling promotes ischemic damage and contributes to edema formation (Colombo et al., 2024). The imbalance in TrkB isoforms induced by *in vivo* injury could be reproduced in a model of *in vitro* excitotoxicity, where it significantly contributed to neuronal death (Vidaurre et al., 2012). This is important because excitotoxicity is not only a mechanism central to stroke but it is also associated to other acute (hypoglycemia, epilepsy, acute trauma) or chronic CNS disorders such as NDDs (Choi, 1988). Three mechanisms induced by excitotoxicity contribute to aberrant TrkB function: (1) modification of the normal ratio of the isoform mRNAs, favoring TrkB-T1 over TrkB-FL mRNA (Gomes et al., 2012; Vidaurre et al., 2012); (2) TrkB-FL cleavage by calpain, resulting in a non-functional truncated receptor similar to TrkB-T1, and a 32 kDa cytosolic fragment containing the TK domain (Tejeda et al., 2016); (3) regulated intermembrane proteolysis (RIP) of both receptor isoforms by sequential action of metalloproteinases (MPs), shedding identical ectodomains acting as BDNF scavengers, and γ-secretases, releasing the isoforms intracellular domains (ICDs) (Tejeda et al., 2016). RIP is a major mechanism of TrkB-T1 regulation both *in vitro* and after ischemia while it is only secondary for TrkB-FL (Tejeda et al., 2016). Similarly to other RIP substrates, TrkB-T1-ICD could have the potential to trigger local signaling at the site of cleavage or modify gene expression after nuclear translocation (Lee and Ch’ng, 2020). Altogether, these pathological mechanisms lead to abnormal BDNF/TrkB signaling which contributes to excitotoxicity, pointing to TrkB-FL and TrkB-T1 as relevant therapeutic targets for treatment of stroke and additional pathologies associated to excitotoxicity. For TrkB-FL, we have designed a cell-penetrating peptide (CPP) able to cross the BBB and plasma membrane that interferes excitotoxicity-induced TrkB-FL interaction with an endosomal protein and retrograde transport to the Golgi complex, secondarily preventing receptor processing (Tejeda et al., 2019) and organelle disruption (Esteban-Ortega and Díaz-Guerra, 2024), an important hallmark of neurodegeneration. In a model of ischemia, this neuroprotective peptide efficiently reduces infarct size and neurological damage (Tejeda et al., 2019).

The importance of TrkB-T1 in the excitotoxic process and the possibility to target both neurons and glial cells strongly support the development of alternative or complementary therapeutic strategies directed to this isoform. In this work, we hypothesize that TrkB-T1 interactions established by isoform-specific C-ter sequences might play a relevant role in neurotoxicity and/or reactive gliosis. Therefore, their interference might prevent those pathological processes and decrease the ischemic damage. Relying again on CPPs, we have designed TrkB-T1-derived peptides containing its C-ter sequence to isolate and identify the isoform protein interactome in basal and excitotoxic conditions. Additionally, we demonstrate that these peptides regulate neuroinflammation and are neuroprotective against excitotoxicity induced *in vitro* or *in vivo,* suggesting that they might be relevant for the therapy of human stroke and other pathologies associated to excitotoxicity and neuroinflammation.

## Results

### Identification and targeting of TrkB-T1-specific interactome in different biological conditions using a CPP containing the isoform C-ter sequence

TrkB-T1 independent functions might require particular protein interactions established by its highly conserved and isoform-specific C-ter (aa 466-476, Figure 1A) (Tessarollo and Yanpallewar, 2022), which could be altered by specific biological conditions. This is the case for RhoGDI1, the only TrkB-T1-interacting protein identified before (Ohira et al., 2005). Additional TrkB-T1-specific interactions established in neurons and astrocytes and how they might be affected by BDNF binding or excitotoxic conditions are presently unknown. To address this, we designed a CPP denominated Bio-sTT1_Ct_ containing three elements: 1) a biotin molecule that allows peptide visualization and pull-down of interacting proteins; 2) a short basic domain from HIV-1 Tat protein, which allows crossing of the BBB and plasma membrane to attached cargoes (Milletti, 2012; Regberg et al., 2013) and 3) the isoform-specific sequence (Figure 1A). As a negative control, we used a similar peptide containing unrelated sequences corresponding to c-Myc (Bio-TMyc). This peptide could enter NeuN+ (arrowheads) and NeuN-cells (asterisk) present in mixed primary cultures of rat embryonic cortex (Figure 1Bd-f), while it showed no effects on neuronal viability in basal or excitotoxic conditions (Ayuso-Dolado et al., 2021; Tejeda et al., 2019). The major cell subtype in these cultures corresponds to neurons which express lower TrkB-T1 levels compared to astrocytes (Figure 1Be). A vesiculated distribution was observed for Bio-sTT1_Ct_ in neurons (Figure 1Bg), resembling that of endogenous TrkB-T1 (Figure 1Bb, e). In fact, double staining with Fluorescein Avidin D and the isoform-specific antibody, prepared with the same TrkB-T1 sequence included in Bio-sTT1_Ct_, suggested that Bio-sTT1_Ct_ and TrkB-T1 mostly colocalized (Figure 1Bi). Next, we confirmed peptide entry into both neurons and astrocytes, as suggested by previous experiment, by performing double staining with cell-type specific markers, respectively NeuN and GFAP (Figure 1C). Bio-sTT1_Ct_ and Bio-TMyc were found inside neurons (arrowheads) and astrocytes (asterisks) alike.

**Figure 1.**
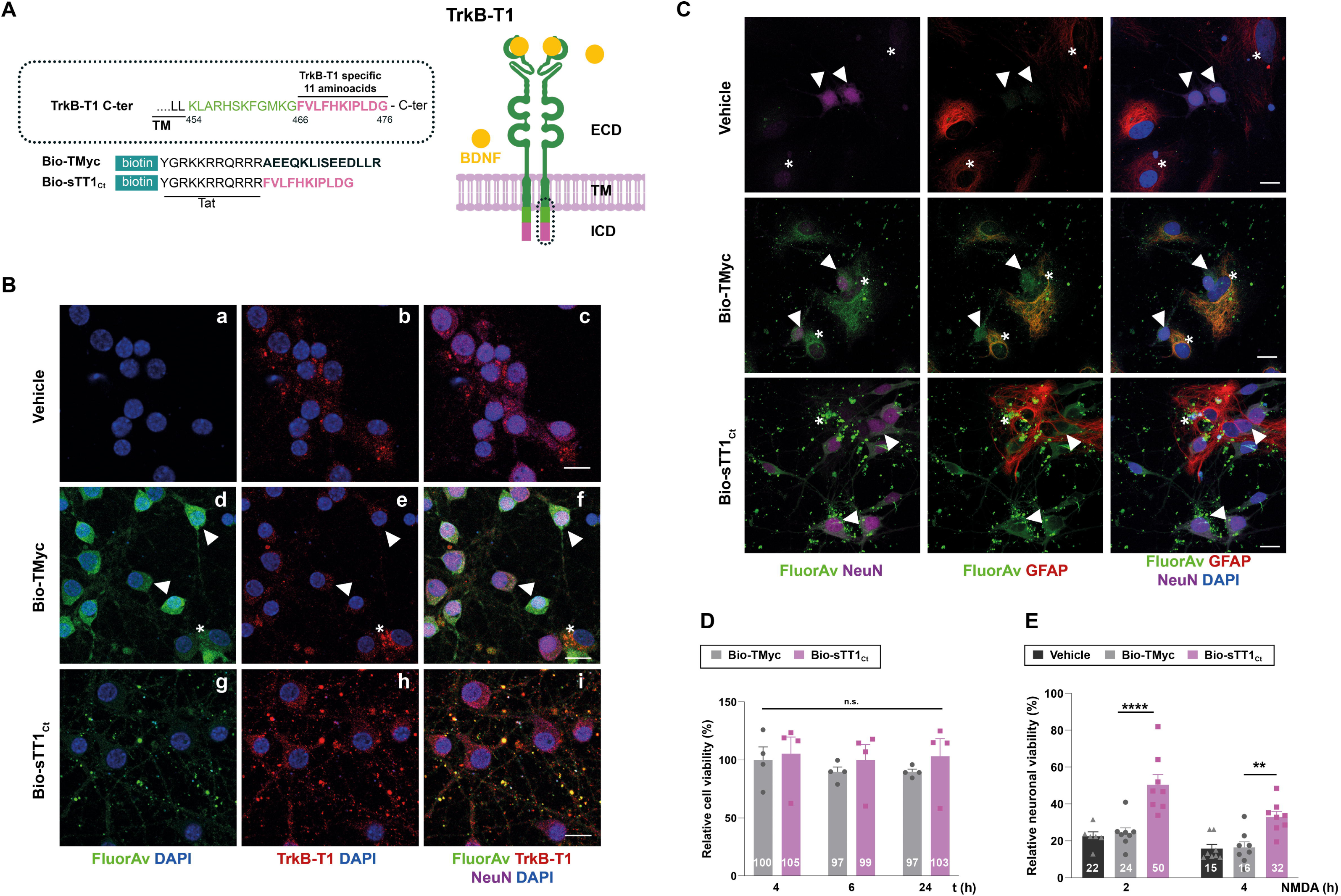
Validation of isoform-specific CPPs as tools for identification of TrkB-T1 interactome and prevention of neuronal death by excitotoxicity. (A) Structure of TrkB-T1 receptor, indicating the extracellular domain (ECD), responsible of brain derived neurotrophic factor (BDNF)-binding, the transmembrane segment (TM) and the short intracellular domain (ICD). The precise sequence corresponding to the TrkB-T1 C-ter (dotted oval) is indicated. It contains a region shared with TrkB-FL (black and green) followed by the TrkB-T1-specific sequence (pink). Biotin (Bio)-labeled Tat-derived CPPs, containing this isoform-specific sequence (Bio-sTT1_Ct_) or unrelated sequences for the control peptide (Bio-TMyc), are also indicated. (B) Immunocytochemistry assays of primary cortical cultures treated with Bio-sTT1_Ct_, Bio-TMyc (25 µM) or vehicle for 30 min. TrkB-T1 and Bio-sTT1_Ct_ distribution were analyzed with an isoform specific antibody (red). Peptide visualization with Fluorescein Avidin D (green) shows that Bio-sTT1_Ct_ presents a pattern similar to that observed for endogenous TrkB-T1. Bio-TMyc distribution was visualized in neurons (arrowheads) and astrocytes (asterisk) (d-f). Scale bar, 10 µm. (C) Analysis of cultures treated as before with specific antibodies for astrocytes (GFAP, red) or neurons (NeuN, magenta). Peptide visualization with Fluorescein Avidin D (green) shows distribution in both neurons (arrowheads) and astrocytes (asterisk). Scale bar, 10 μm. (D) Cell viability of cortical cultures treated with Bio-sTT1_Ct_ and Bio-TMyc (25 µM) for 4, 6 or 24 h. Means ± SEM and individual points are presented relative to values obtained for 4 h of Bio-TMyc treatment (100%). Data were analyzed using two-way ANOVA test followed by *post hoc* Bonferroni test, n = 4. (E) Neuronal viability in cultures incubated with Bio-TMyc or Bio-sTT1_Ct_ (25 µM) for 30 min and treated with NMDA for 2 or 4 h. Means ± SEM and individual points are presented relative to the values obtained for untreated cells (100%). Data were analyzed using two-way ANOVA test followed by *post hoc* Bonferroni test, n = 8.

Above data suggested that Bio-sTT1_Ct_ might interfere protein interactions established by TrkB-T1 C-ter, for example those required for promotion of neuronal death in a context of excitotoxicity. The incubation of primary cortical cultures with Bio-TMyc or Bio-sTT1_Ct_ (4-24 h) showed no toxicity in basal conditions (Figure 1D). The induction of excitotoxicity by overactivation of NMDARs with co-agonists NMDA (100 µM) and glycine (10 µM), herein simply denoted as NMDA, dramatically decreased neuronal viability in a time-dependent manner in the absence of peptide or after preincubation with control peptide Bio-TMyc (Figure 1E). In contrast, Bio-sTT1_Ct_ presented a significant neuroprotective effect after 2 (50 ± 5%) or 4 h (32 ± 3%) of NMDA treatment compared to those obtained in cultures incubated with Bio-TMyc (24 ± 3%, \*\*\*\**p*<0.0001, for 2 h; 16 ± 3%, \*\**p*<0.01, for 4 h; n = 8).

We interpreted these results as indicative that peptide Bio-sTT1_Ct_ was indeed able to interfere protein interactions established by TrkB-T1 inside cells and, thus, could be used to isolate and identify this isoform C-ter interactome in different biological conditions. Therefore, cultures were incubated with Bio-TMyc or Bio-sTT1_Ct_ (30 min) before treatment with BDNF (100 ng/ml) or NMDA (100 µM) for 30 additional minutes or left untreated (Figure 2A). Cell lysis was performed in conditions that do not disrupt peptide-protein interactions and then, complexes formed inside cells by the biotin-labeled peptides were isolated using streptavidin-agarose precipitation. After that, proteins interacting with Bio-TMyc and Bio-sTT1_Ct_ were identified by a label free proteomic assay, obtaining a total of 4697 proteins for the three different biological conditions and four independent experiments analyzed. Some proteins might be interacting with shared peptide elements such as the biotin moiety or the Tat sequence, or even streptavidin-agarose (Figure 2A). Therefore, we first evaluated by a Principal Component Analysis (PCA) whether the interactome profiles obtained for Bio-TMyc and Bio-sTT1_Ct_ were different (Figure 2B). The first and second principal components of this analysis (respectively, PC1 and PC2) split the samples into two groups, proving a differential profile of interactions for Bio-TMyc and Bio-sTT1_Ct_.

**Figure 2.**
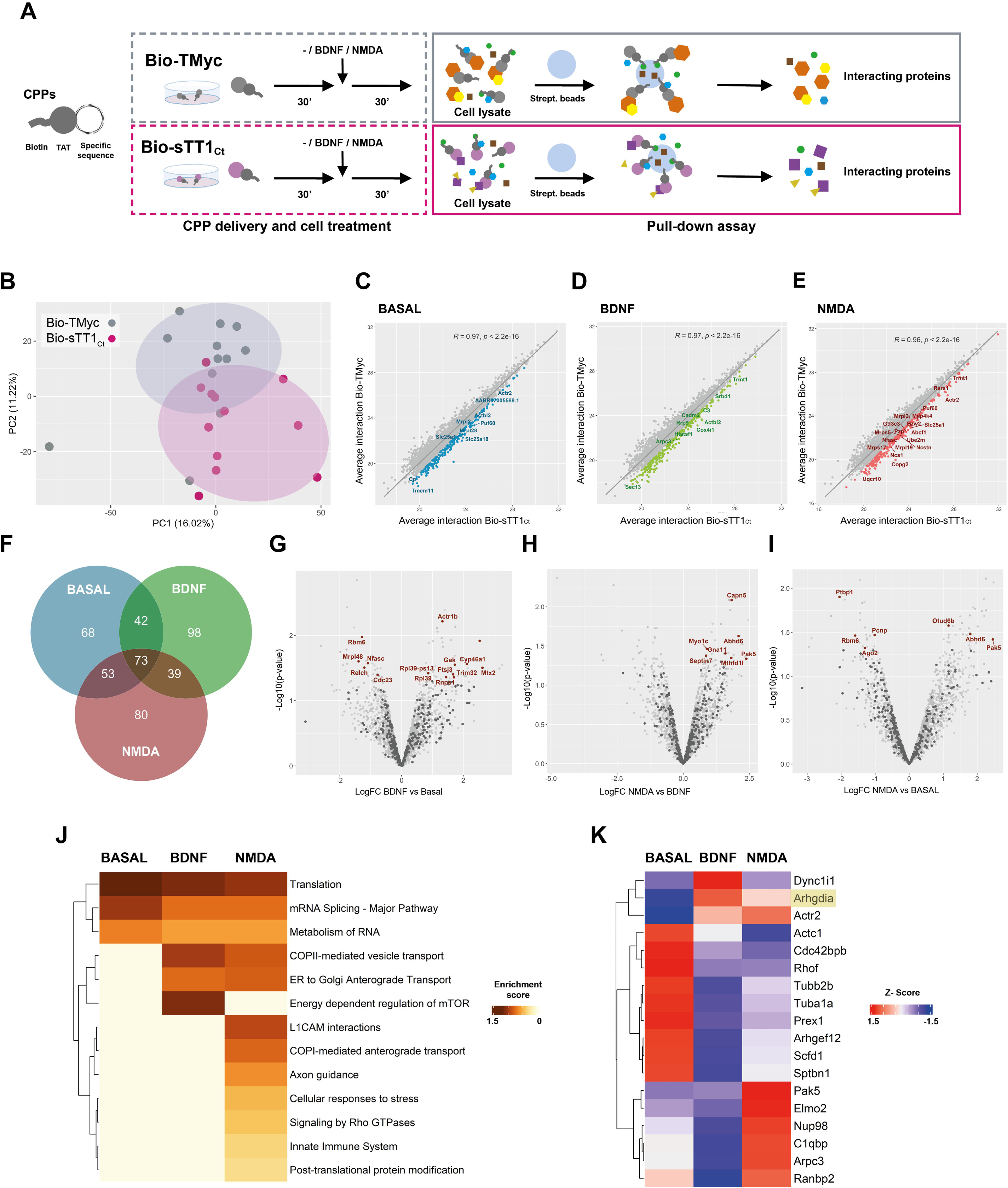
Bio-sTT1_Ct_ as a tool to approach TrkB-T1 specific interactome in different biological conditions. (A) Experimental design of pull-down assays to isolate Bio-TMyc and Bio-sTT1_Ct_ interacting proteins. Cultures were incubated with Bio-sTT1_Ct_ or Bio-TMyc (25 µM) for 30 min before treatment with BDNF (100 ng/mL) or NMDA for 30 min. After that, cell lysates were combined with streptavidin agarose beads to isolate the CPP-interacting proteins. (B) Principal Component Analysis of Bio-TMyc and Bio-sTT1_Ct_ pull-down isolates. Samples are represented using the first (PC1) and second (PC2) components of the analysis. (C-E) Pearson’s correlation of the average protein interactions established by Bio-TMyc and Bio-sTT1_Ct_ in basal conditions (C), or after BDNF (D) or NMDA treatment (E). Colored points represent the top 20% proteins whose residuals are furthest from the correlation line and have higher levels of binding to Bio-sTT1_Ct_. (F) Venn diagram representing the number of proteins selected for each condition following the criteria described above. (G-I) Volcano plot presenting the results of the differential analysis of interacting proteins comparing BDNF vs basal conditions (G), NMDA vs BDNF (H) and NMDA vs basal conditions (I). Log_2_FC and -Log_10_(p-value) are shown. Proteins selected with the criteria explained above are represented as dark gray points while proteins showing statistically significant differences are labeled and presented as brown dots (p-value < 0.05). FDR = 9.52% (G), 4.15%) (H) or 9.55% (I). FC, fold change. (J) Pathway enrichment analysis of selected proteins for basal, BDNF and NMDA conditions. A heatmap showing the enrichment score for each pathway (Reactome.db) at the different conditions is presented. (K) Comparison of RhoGDI (*Arhgdia*) (highlighted) and our selected proteins annotated in “Signaling by Rho GTPases” Reactome pathway in basal, BDNF and NMDA conditions. A heatmap showing the z-score indicating levels for each protein at the different conditions is presented.

To bring into focus those proteins more specifically interacting with Bio-sTT1_Ct_, we calculated Pearson’s correlation between the average protein interactions established by Bio-TMyc and Bio-sTT1_Ct_ at the different conditions (Figure 2C-2E). Then, we analyzed the correlation line residuals and selected the top 20% of proteins having higher average interaction with Bio-sTT1_Ct_ compared to Bio-TMyc (highlighted as blue, green or red dots, respectively for the basal, BDNF or NMDA conditions). Some of the 453 selected proteins maintained Bio-sTT1Ct-interaction at all three conditions while others were affected by BDNF or NMDA treatment (Figure 2F). Interestingly, the gene ontology (GO) analysis of biological process enrichment (string.db) associated with Bio-sTT1_Ct_-interacting proteins presented terms related to gene expression and mRNA regulation, including splicing, at all treatment conditions (Figure S1). Only upon NMDA treatment, other biological process terms appeared such as cytoskeletal reorganization and cell division (Figure S1C). Altogether, these results suggested a possible role of protein complexes established by TrkB-T1 C-ter in cellular processes relevant to gene expression and, in primary cultures subjected to excitotoxicity, cytoskeletal organization and cell division.

Then, we further characterized the impact on Bio-sTT1_Ct_ interactions of different cellular conditions by differential analysis of the binding observed for the selected group of proteins in basal conditions or after BDNF or NMDA treatment (Figure 2G-2I). Significant differences were found in all comparisons (named proteins in corresponding figures, summarized in Tables S1-S3), proving that some of the protein complexes formed with Bio-sTT1_Ct_ were affected by the applied biological stimuli. Thus, for example, BDNF induced Bio-sTT1_Ct_-binding of proteins involved in endocytosis and molecular trafficking (*Gak*), cholesterol metabolism (*CYP46A1*), pre-rRNA processing (*Ftsj3*) or mitochondria-endoplasmic reticulum contact sites (MERCS) (*Mtx2/Grp75*, Figure 2G). Meanwhile, excitotoxicity promoted significant changes in binding of proteins involved, among others, in gene expression (*Otud6b*, increased; *Ago2*, *Rbm6* and *Ptbp1*, decreased; Figure 2I) and cytoskeletal remodeling (*Septin7*, *Pak5*, *Capn5*; Figure 2H and 2I). Furthermore, when we compared the enriched pathways (Reactome, string.db) among the three tested conditions (Figure 2J), we observed that peptide Bio-sTT1_Ct_ was interacting in all of them with proteins involved in translation, RNA metabolism and splicing, as seen before in the biological process terms (Figure S1), while BDNF treatment enriched pathways related to vesicle transport and mTOR regulation. In fact, vesicle transport was also increased with NMDA, while other pathways key for the promotion of the neurotoxic process such as innate immune system, L1CAM interactions, cellular response to stress, axon guidance and RhoGTPases activity appeared. These changes suggest an important role of TrkB-T1 C-ter sequence in the development of the excitotoxic process by interaction with proteins that would be involved in this pathological process.

Next, we analyzed in detail the group of selected Bio-sTT1_Ct_-interacting proteins included in the Reactome function “signaling by RhoGTPases” (Table S4 and Figure 2K), a pathway enriched after NMDA treatment (Figure 2J). This analysis was important because this family of proteins is central to neuronal survival and neurodegeneration in addition to neuronal development (Stankiewicz and Linseman, 2014). Although specific Bio-sTT1_Ct_-binding to RhoGDI was low in the cortical cultures and this protein did not pass the established selection criteria, we decided to include it in this analysis because RhoGDI is a TrkB-T1-interacting protein in astrocytes whose binding is negatively regulated by BDNF (Ohira et al., 2005). In contrast, binding of RhoGDI (*Arhgdia* encoded) to Bio-sTT1_Ct_ in the untreated neuronal-enriched cultures was very low compared to that of other RhoGTPase family members and increased after BDNF and NMDA treatment (Figure 2K), although total RhoGDI levels were not affected by these treatments (Figure S2). Thus, TrkB-T1/RhoGDI interaction might be different in neurons and astrocytes. Regarding other RhoGTPase-related proteins (Table S4), most of them presented a very reduced binding after BDNF stimulation while two opposite Bio-sTT1_Ct_-interacting patterns were found in excitotoxicity compared to the basal conditions. Thus, the interaction was downregulated by NMDA treatment for an important group of proteins, including cytoskeleton components (*Actc1*, *Tubb2b*, *Tuba1a*, *Sptbn1*), Rho and Rac GEFs (respectively *Arhgef12* and *Prex1*) or a Cdc42 effector (*Cdc42bpb*). In contrast, Bio-sTT1_Ct_-interaction was promoted by NMDA for some proteins including *Actr2* and *Arpc3,* components of the Arp2/3 complex mediating actin polymerization, *C1qbp*, involved in inflammation, *Nup98* and *Ranbp2,* components of the nuclear pore complex (NPC) and involved in RNA transport, *Elmo2*, a regulator of Rac GTPase activity and *Pak5*, a downstream effector of Rac1 and Cdc42 GTPases.

Altogether, these results represent a first comprehensive analysis of the protein interactions established by TrkB-T1 isoform-specific C-ter sequence, and how they are modulated in cell models of physiological and pathological conditions. Some of the identified TrkB-T1-interacting proteins might contribute to the central role played by this isoform in excitotoxicity and ischemia and help to explain preliminary results showing a neuroprotective effect of peptide Bio-sTT1_Ct_ (Figure 1E), unveiling a new therapeutic strategy to treat stroke and additional pathologies associated to excitotoxicity that merits further investigation.

### Design of peptide TT1_Ct_ and mechanism of neuroprotection in cellular models of excitotoxicity

To further characterize the mechanism of action and effects of TrkB-T1 interference in models of excitotoxicity, we designed a new CPP, designated as TT1_Ct_ (Figure 3A). This peptide lacks the biotin molecule, to avoid possible unwanted effects, and contains two spacer prolines separating Tat and isoform-specific sequences, to disrupt a possible regular secondary structure between these functionally-independent moieties that might affect peptide stability and efficacy. As a negative control, we used TMyc, a similar peptide having unrelated c-Myc sequences (Figure 3A) (Ayuso-Dolado et al., 2021; Tejeda et al., 2019). We first analyzed if TT1_Ct_ had toxic effects on neurons under basal conditions as well as neuroprotective effects after induction of *in vitro* excitotoxicity. We measured neuronal viability in a dose-response experiment using different TMyc and TT1_Ct_ (5, 15 or 25 µM) concentrations for 2.5 h (Figure 3B). In the basal conditions, TT1_Ct_ was toxic at the higher dose employed, probably by mimicking some TrkB-T1-ICD neurotoxic effect, indicating the importance of CPP dosage for future experiments. Accordingly, TT1_Ct_ preincubation (25 µM, 30 min) followed by a chronic NMDA treatment (2 h) was not neuroprotective and neuronal viability decreased to values similar to those obtained with the control peptide. In contrast, we observed a strong TT1_Ct_ neuroprotective effect for peptide concentrations of 5 or 15 µM that did not promote neuronal death in the basal conditions (Figure 3B). After NMDA treatment, neuronal viability reached values of 46 ± 7% for 15 µM TT1_Ct_, significantly higher than those observed in cultures incubated with the same concentration of TMyc (13 ± 9%, \*\**p*<0.01; n = 8). Next, we compared the neuroprotective efficacy of TrkB-T1-derived peptides, Bio-sTT1_Ct_ and TT1_Ct_ (15 µM), in a time-course experiment (Figure 3C). In cultures preincubated as before and subjected to excitotoxicity for 2 or 4 h, similar neuroprotective effects were obtained for both peptides. Neuroprotection due to TT1_Ct_ preincubation was maintained when the peptide was applied at the time of damage induction but lost if added at later times (Figure S3), suggesting that this peptide affects processes taking place in cultures at early times of NMDAR overactivation.

**Figure 3.**
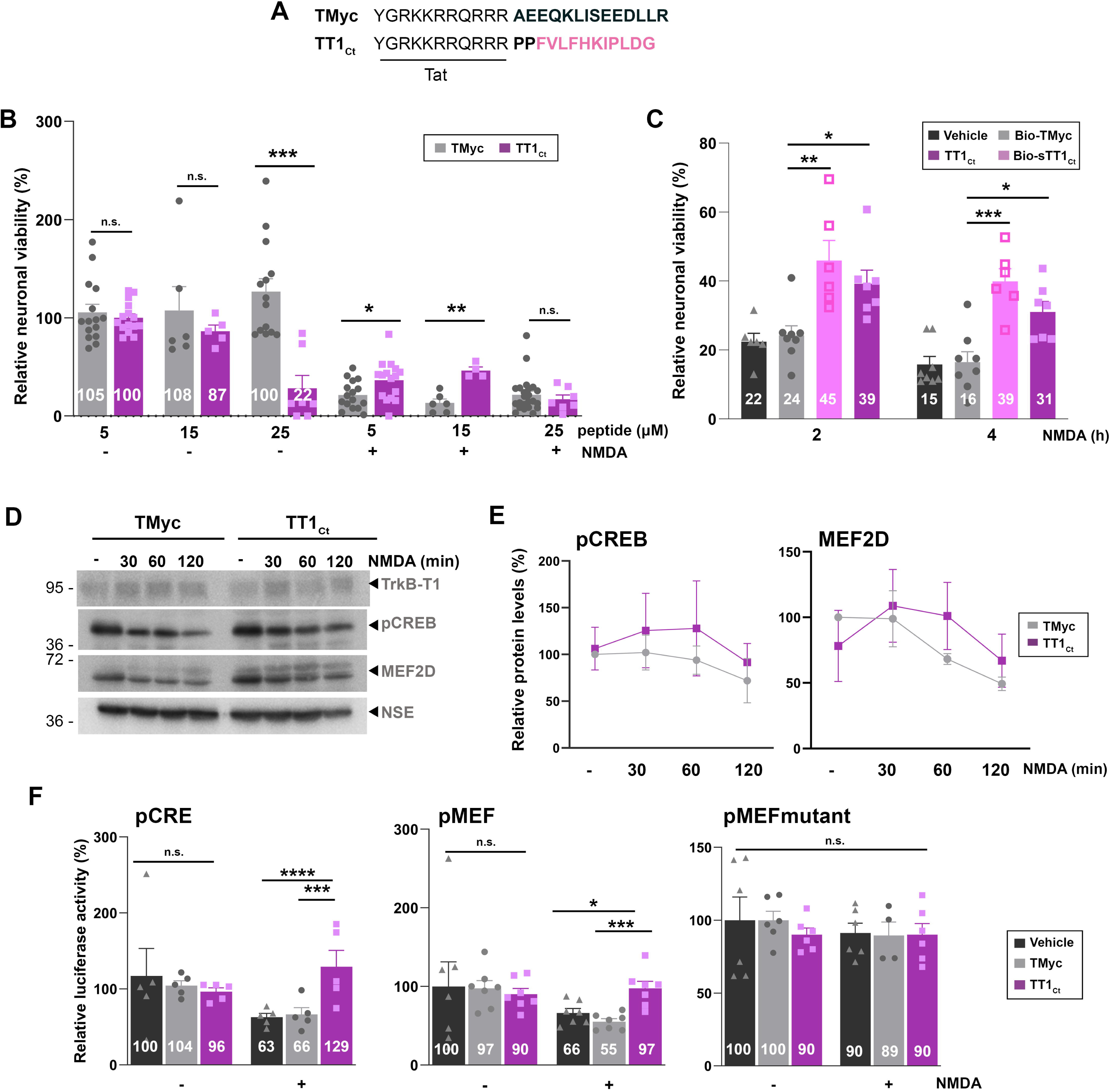
TrkB-T1-derived peptide TT1_Ct_ is neuroprotective against *in vitro* excitotoxicity and prevents a decrease in CRE and MEF promoter activities induced by the excitotoxic injury. (A) Sequence of control peptide (TMyc) and TrkB-T1-derived CPP (TT1_Ct_). (B) Neuronal viability in cultures incubated with TMyc or TT1_Ct_ (5, 15 or 25 µM) for 30 min and then treated with NMDA (100 µM) for 4 h or left untreated. Individual data and means ± SEM are presented relative to values obtained in the untreated cells (100%). Data were analyzed using Kruskal-Wallis test followed by Mann-Whitney U-test, n = 6-16. (C) Neuronal viability in cultures incubated with Bio-TMyc, Bio-sTT1_Ct_ or TT1_Ct_ (15 µM) for 30 min and then treated with NMDA (100 µM) for 2 or 4 h. Individual data and means ± SEM are presented relative to values obtained for untreated cells (100%). Data were analyzed using two-way ANOVA test followed by *post hoc* Bonferroni test, n = 6-8. (D) Western Blot analysis of TrkB-T1, pCREB and MEF2D levels in cultures incubated with TMyc or TT1_Ct_ (15 µM) for 30 min followed by treatment for 30, 60 or 120 min with NMDA. A representative experiment is shown. (E) Quantitation by densitometric analysis of pCREB and MEF2D levels. Means ± SEM are presented relative to basal conditions in the presence of TMyc (100%), n = 5. (F) Effect of excitotoxicity and TT1_Ct_ treatment on CRE and MEF2 promoter activity. Cultures transfected with plasmids containing minimal CREB or MEF2 response elements (respectively, pCRE and pMEF) or pMEFmut were preincubated with peptides as above and treated with NMDA for 2 h or left untreated. Individual results and means ± SEM are presented relative to luciferase expression in untreated cultures. Data was analyzed by two-way ANOVA test followed by *post hoc* Bonferroni test, n = 5-7. For cultures transfected with pCRE and pMEF, treated with vehicle or TMyc, differences between -/+ NMDA are statistically significant although not shown for simplicity.

Our next goal was to investigate the mechanism of neuroprotection by TT1_Ct_ action. We have shown before that excitotoxicity induces a progressive increase in TrkB-T1 levels (Vidaurre et al., 2012) which probably contributes to excitotoxic neuronal death. Other proteins also affected by NMDAR overactivation are prosurvival TFs CREB and MEF2. Levels of S133 phosphorylated CREB (pCREB), generally considered the active protein, are strongly reduced early upon induction of excitotoxicity probably due to phosphatase activation. The result is CREB shut-off, inhibition of BDNF expression and death of mature neurons (Hardingham et al., 2002). In addition, there is a severe decrease of MEF2D in cultures subjected to excitotoxicity (Tejeda et al., 2019), probably due to action of caspases or calpain (Tang et al., 2005; Wei et al., 2012). We observed that the effect of excitotoxicity in cultures incubated with TMyc (15 µM, 30 min) before NMDA treatment (0-120 min) was as expected (Figures 3D and 3E). At these early times of excitotoxicity, preincubation with TT1_Ct_ did not have significant effects in TrkB-T1 regulation (Figure 3D). In contrast, this peptide was able to partially preserve pCREB and MEF2D levels, although these differences were not statistically significant compared to TMyc-treated cultures (Figure 3E). In these experiments, neuron-specific enolase (NSE), a protein not affected by NMDA, was used as a loading control and for protein normalization. Altogether, these results suggest that one mechanism of TT1_Ct_ neuroprotection might be mediated by TFs CREB and MEF2D through regulation of gene expression. To further explore the potential role of TT1_Ct_ on CREB and MEF2, we performed reporter assays using promoters with minimal response elements (respectively, pMEF2 or pCRE; Woronicz et al, 1995; Deogracias et al, 2004; see Key Resources Table) regulating luciferase expression. In basal conditions, incubation with TT1_Ct_ (15 µM) for 2.5 h had no significant effect on pCRE and pMEF promoter activities compared to cultures treated with the same concentration of TMyc or without peptide (Figure 3F). In contrast, induction of excitotoxicity provoked a dramatic decrease in luciferase activity when cultures were preincubated with TMyc or vehicle. However, the reduction in CREB and MEF2 promoter activities was significantly prevented in the presence of TT1_Ct_. An inactive pMEF mutant with reduced luciferase expression was unresponsive to NMDA or peptide treatment, proving that excitotoxicity regulates MEF2-promoter activity and TT1_Ct_ has a specific effect on it.

Above results are important because these TFs are central to neuronal survival induced by synaptic activity (Linseman et al., 2003; Lonze and Ginty, 2002) or neurotrophins (Bonni et al., 1999; Liu et al., 2003). In addition, CREB also regulates the activity of astrocytes in response to neurotransmitters (Carriba et al., 2012; Karki et al., 2013; Modi et al., 2015). To establish if the protective role of TT1_Ct_ alters the excitotoxicity-induced transcriptional changes in neurons and/or astrocytes, we next analyzed the levels of mRNAs encoding several proteins involved in survival/death choices (Figure 4). As expected, in the presence of TMyc, NMDA induced a strong decrease in levels of mRNAs encoding GluN1 (Gascon et al., 2005), GluN2A (Gascon et al., 2005) and TrkB-FL (Vidaurre et al., 2012) which did not respond to preincubation with TT1_Ct_ (Figures 4A-4C). In contrast, we confirmed an increase in TrkB-T1 mRNA induced by NMDA (Gascon et al., 2005) in the presence of TMyc, that was counteracted by TT1_Ct_ action (Figure 4D). For BDNF, excitotoxicity showed a tendency to increase mRNA levels in the presence of TMyc, in agreement to previous results (Zafra et al, 1991), and such accumulation was strongly exacerbated by TT1_Ct_ (Figure 4E). Finally, this peptide decreased GFAP mRNA levels relative to those found in TMyc-treated cultures in excitotoxicity (Figure 4F). In conclusion, for the analyzed mRNAs, TT1_Ct_ does not seem to affect the transcriptional downregulation of neuronal genes induced by excitotoxicity while it strongly modifies the expression of genes expressed in astrocytes (GFAP) or both neurons and astrocytes (TrkB-T1 and BDNF), where it counteracts the effects of excitotoxicity. Altogether, we conclude that interference of TrkB-T1 protein interactions by TT1_Ct_ during excitotoxicity maintains levels and promoter activities of CREB and MEF2, which regulate the transcription of TrkB-T1 itself and, other target genes critical to neuronal survival and astrocyte activation.

**Figure 4.**
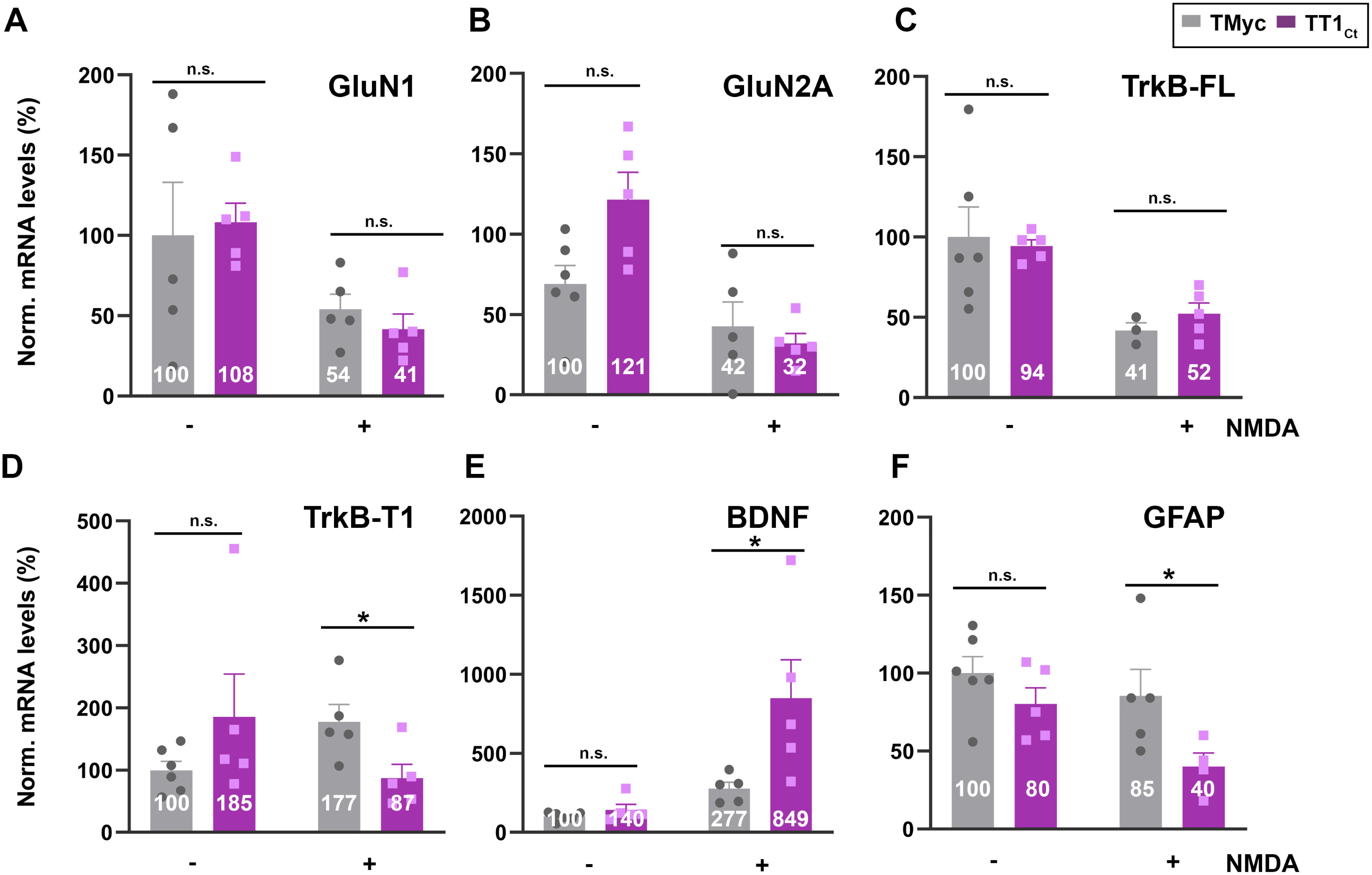
TT1_Ct_ interferes transcriptional changes induced by excitotoxicity affecting expression of genes involved in survival/death choices. Cultures preincubated with TT1_Ct_ or TMyc (15 µM) for 30 min were treated with NMDA for 4 h or left untreated. Levels of mRNAs encoding for NMDAR-subunits GluN1 (A) and GluN2A (B) or TrkB-FL (C) and TrkB-T1 isoforms (D) were normalized to those of NSE. Levels of BDNF (E) and GFAP mRNA (F) were normalized relative to GAPDH. Individual values and means ± SEM are presented relative to gene expression in cultures treated with TMyc (100%). Data were analyzed by two-way ANOVA test followed by *post hoc* Tukey’s or Kruskal Wallis test followed by *post hoc* Dunn’s test, n = 5.

### Effects of TT1_Ct_ in a model of ischemic stroke induced by photothrombosis

Our next objective was to investigate if TT1_Ct_ could counteract brain degeneration and interfere with TrkB-T1/GFAP expression in ischemic stroke, where the process of excitotoxicity takes place *in vivo*. First, we demonstrated that the biotinylated forms of TT1_Ct_ (Bio-TT1_Ct_) and Bio-TMyc, intravenous (i.v.) injected into control animals for 30 min, were able to cross the BBB and efficiently reached most cortical cells and distributed in cell bodies (asterisks) and projections (arrowheads) (Figure S4). Then, we selected a mouse model of permanent ischemia induced by microvascular photothrombosis, which models the embolic or thrombotic occlusion of small arteries often produced in human stroke (Pevsner et al., 2001). Photothrombosis causes early breaking of the BBB (Abdullahi et al., 2018) and damage of cortical motor and somatosensory regions, with infarcts detected after 3 h (Tejeda et al., 2019) and reaching their maximum volumes by 24 h (Figure 5A). In animals sacrificed 5 h after damage induction, we confirmed a strong decrease of TrkB-FL in neurons of the infarcted area, identified by condensed nuclei indicative of cell injury (Figure 5B). In contrast, levels of TrkB-T1 and GFAP already increased in the infarcted area at this early time of damage, particularly in cells located in the interface between ischemic and non-ischemic tissue which might correspond to the emerging glial scar. Similar results were obtained in animals injected with peptide Bio-TMyc (3 nmol/g, i.v.) 1 h after damage initiation (Figure S5). Next, we analyzed Bio-TMyc and Bio-TT1_Ct_ distribution in the cortex of 5 h post-stroke male and female mice, detecting the tagged peptides in both neuronal (arrows), and non-neuronal cells (asterisks) of the contralateral hemisphere (Figure 5C). Detection of the biotinylated peptides at least 4 h after administration suggests relative peptide stability in brain tissue.

**Figure 5.**
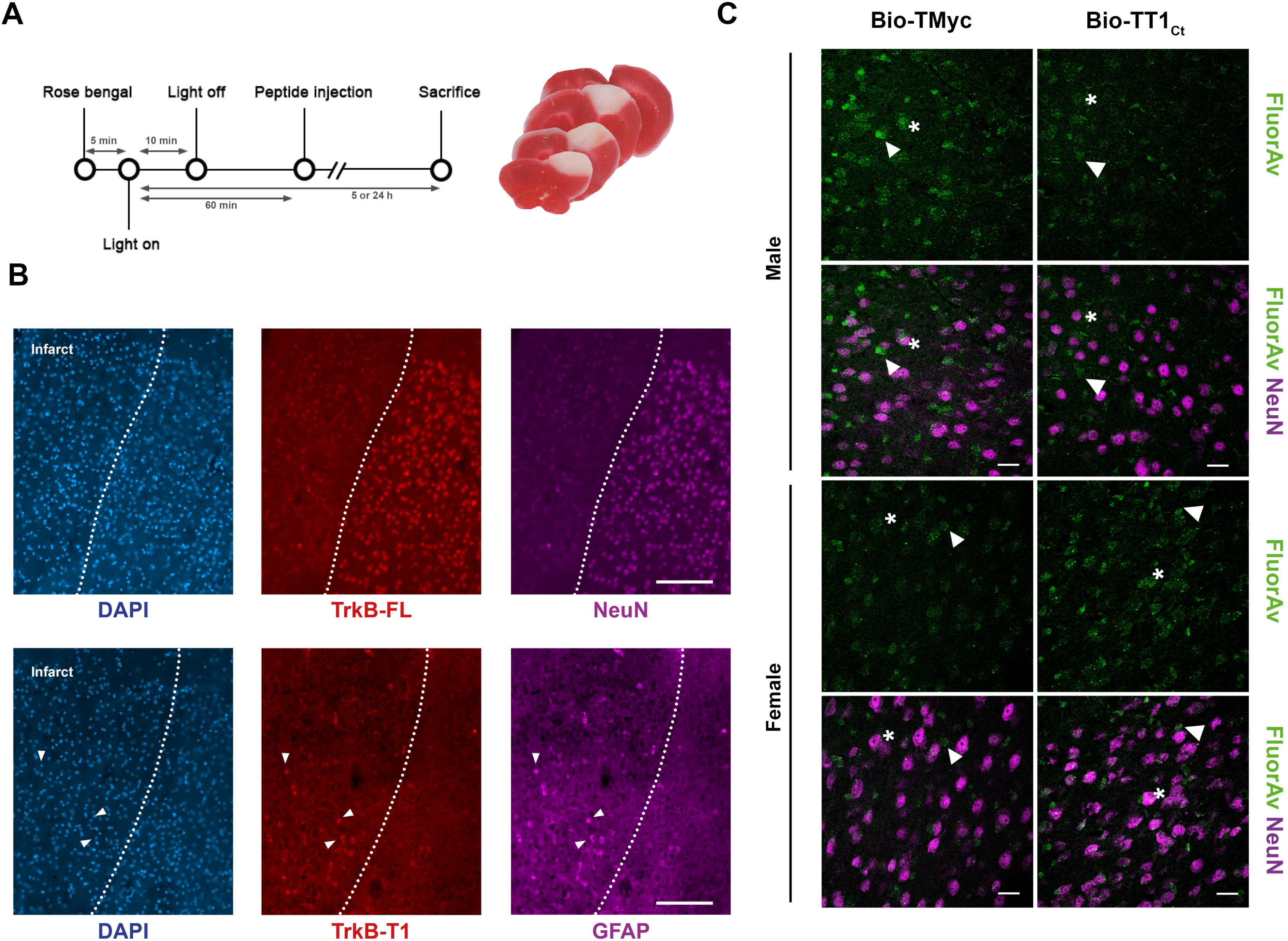
TT1_Ct_ brain distribution in a mouse model of permanent ischemia showing upregulation of TrkB-T1+/GFAP+ cells in the interface between ischemic and non-ischemic tissue. (A) Experimental design to analyze *in vivo* effects of TMyc and TT1_Ct_. Permanent vessel occlusion and focal brain damage were induced in mice by cold-light irradiation after Rose Bengal i.v. injection. CPPs (3 nmol/g) were i.v. injected 1 h after damage initiation. Animals were sacrificed 5 or 24 h after injury onset as indicated. Representative 1 mm brain coronal slices stained with TTC after 24 h of insult are shown. (B) Immunohistochemistry of brain coronal sections prepared from male animals 5 h after insult stained with isoform-specific TrkB antibodies (TrkB-FL and TrkB-T1), NeuN, GFAP and DAPI. Representative cell observer images of cortical areas of the infarct border are shown. Arrows indicate concurrent increased expression of TrkB-T1 and GFAP in astrocytes. Scale bar, 50 μm. (C) Analysis of TT1_Ct_ delivery to male and female mice cortex after 5 h of ischemia. Bio-TT1_Ct_ and Bio-TMyc were injected as before and detected in the contralateral region of coronal sections by Fluorescein Avidin D (green). Neuronal marker NeuN (magenta) is also shown. Peptide delivery was observed in neuronal (asterisks) and non-neuronal cells (arrowheads). Representative confocal microscopy images of cortical areas correspond to single sections. Scale bar, 20 μm.

To investigate the effect of the biotinylated peptides in glial subpopulations during ischemic stroke, we first evaluated co-staining of GFAP-labelled astrocytes and complement component 3 (C3) in animals subjected to 5 h (Figure 6A) or 24 h (Figure 6B) of injury. C3 is a marker for inflammatory reactive astrocytes which, among other features, lose the capacity to support neuronal survival and synaptogenesis, and induce death of neurons and mature differentiated oligodendrocytes (Liddelow et al., 2017). Animals treated with TMyc showed most GFAP+ cells stained by the C3 antibody at the infarct boundary, both at 5 and 24 h of damage. Moreover, the number of cells identified as GFAP+/C3+ strongly increased over time. In contrast, since early stages of injury, peptide TT1_Ct_ caused a strong decrease in GFAP+/C3+ cells. In addition to an early TT1_Ct_ effect on glial scar formation, we investigated a possible influence of this peptide on macrophage infiltration and microglia activation in the injured brain (Figures 6C-6G). Protein CD68 (ED1), expressed in macrophages, activated microglia and other cell types, was detected in the infarct border and core of animals treated with the control peptide, while it was absent of the contralateral region (Figure 6C). Most CD68+ cells had a rounded morphology compatible with macrophages, although more elongated cells were also detected (Figure 6D). Treatment with TT1_Ct_ greatly decreased number of CD68+ cells in the infarcted and surrounding tissues, in parallel with reduced immunoglobulin infiltration (Figure S6), which can be detected as early as 2.5 h after damage initiation due to BBB damage (Ayuso-Dolado et al., 2021). Expression of Iba1, a protein present in microglia and circulating macrophages (Figure 6E), was detected in cells with ramified morphology characteristic of resting microglia located in the contralateral region of animals treated with TMyc and TT1_Ct_ (Figure 6E and 6F). Upregulation of Iba1 was observed in the ischemic brain of animals treated with TMyc accompanied by morphological cellular changes, indicative of hypertrophic activated microglia. On the other hand, treatment with TT1_Ct_ limited the upregulation of Iba1 in the reactive microglia and decreased presence of reactive microglia in the damaged tissue (Figure 6E). Double immunostaining with anti-CD68 and Iba1 antibodies showed very little signal overlapping (Figure 6G). Altogether, these results show the importance of TrkB-T1 in glial scar formation and the inflammatory response after an ischemic damage, as well as the possibility to restrain these processes by using the brain-accessible peptide TT1_Ct_.

**Figure 6.**
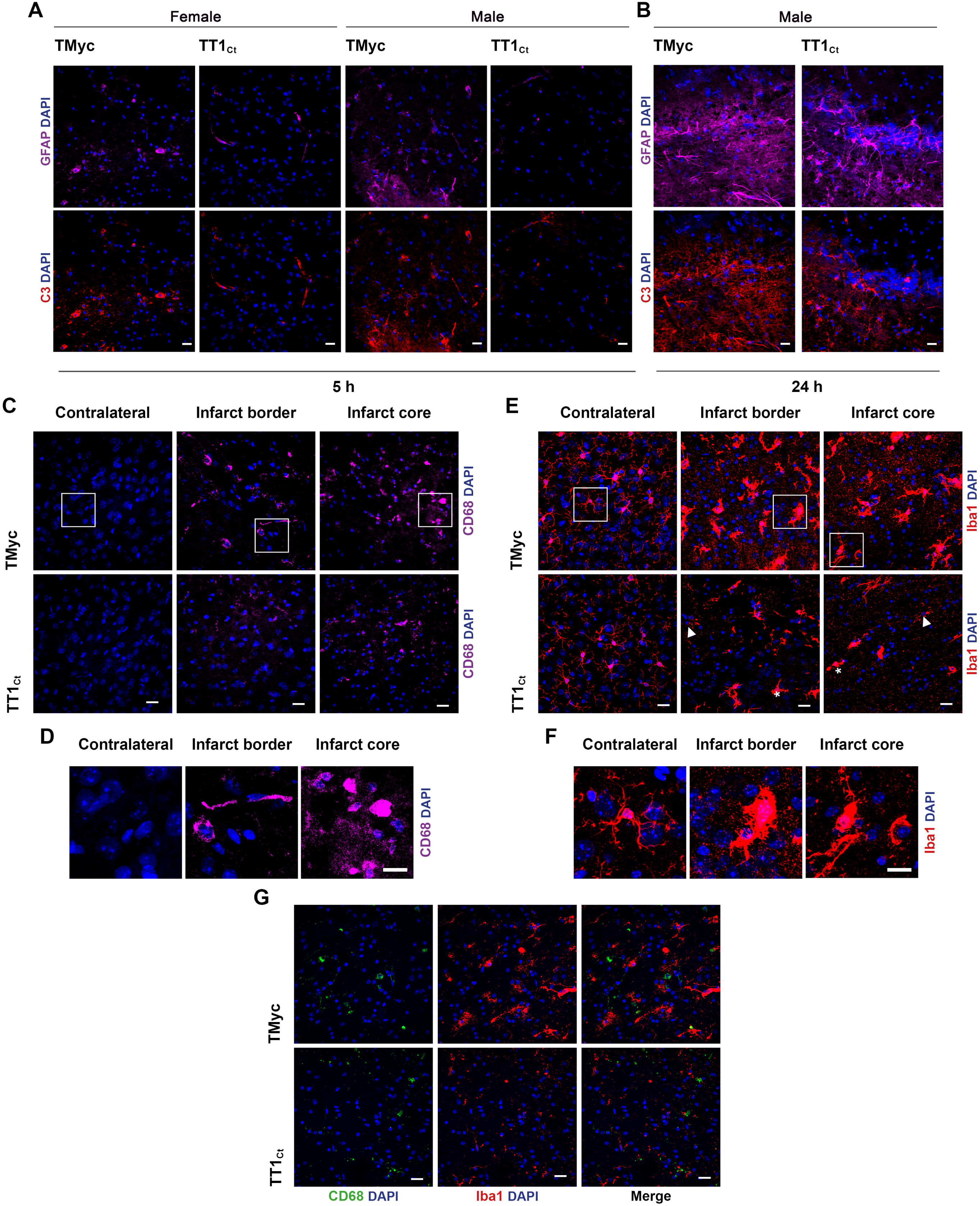
Treatment with TT1_Ct_ prevents reactive gliosis at early and late times of ischemic damage. (A) Coronal sections of male and female mice injected with TMyc or TT1_Ct_ (3 nmol/g) after 5 h of insult were stained with antibodies for C3 (red), GFAP (magenta) and DAPI to detect reactive astrocytes. Representative maximum intensity projection confocal images from the infarct border are shown. Scale bar, 20 μm. (B) Coronal sections of male mice treated with TMyc or TT1_Ct_ (3 nmol/g) after 24 h of insult were stained as above. Representative maximum intensity projection of confocal images from the infarct border are shown. Scale bar, 20 μm. (C) Coronal sections of male mice injected with TMyc or TT1_Ct_ (3 nmol/g) after 24 h of insult were stained with an antibody for CD68 (magenta) and DAPI to detect microglia and macrophage inflammatory state. Representative maximum intensity projection of confocal images from the contralateral cortex, infarct core and infarct border are shown. Scale bar, 20 μm. (D) Detail of cell morphology in selected areas indicated in panel C, corresponding to animals injected with TMyc as above indicated. Scale bar, 20 μm. (E) Coronal sections of male mice injected with TMyc or TT1_Ct_ (3 nmol/g) after 24 h of insult were stained with antibody for Iba1 (red) and DAPI to detect microglia and macrophage inflammatory state. Representative maximum intensity projection of confocal images of the contralateral cortex, infarct core and infarct border are shown. Reactive microglia (asterisks) and resting microglia (arrows) were detected. Scale bar, 20 μm. (F) Detail of cell morphology in selected areas indicated in panel E, corresponding to animals injected with TMyc as above indicated. Scale bar, 20 μm. (G)Double immunohistochemistry with CD68 (green) and Iba1 (red) antibodies of the infarct core of male mice injected with TMyc or TT1_Ct_ as above to analyze possible signal overlapping. Representative maximum intensity projection of confocal images are shown. Scale bar, 20 μm.

Finally, we wanted to establish if the described TT1_Ct_ effects on reactive gliosis resulted in brain protection after ischemic injury and improved the neurological outcome of affected animals. First, we analyzed the effect of TT1_Ct_, administered in both male and female animals 1 h after damage induction, on infarct volume (Figure 7A) and neurological damage (Figure 7B) evaluated at 24 h. This experimental setting mimics a clinical situation where the neuroprotective molecules are provided after insult onset. Treatment with TT1_Ct_ (3 nmol/g, i.v.) significantly reduced the infarct size (10 ± 1% relative to total hemisphere volume), approximately 40% when compared to TMyc treated mice (17 ± 2%, \*\*\**p*<0.001, n = 19; Figure 7A). Curiously, administration of TT1_Ct_ 10 min after damage initiation showed a comparatively modest effect both on infarct volume (23%) and neurological damage (29%) (data not shown). A gender disaggregated data analysis showed, for TMyc-treated animals, a tendency to reduced infarct volume in female mice compared to male (15 ± 2% versus 20 ± 1%; *p =* 0.08, n = 9: Figure 7C). However, treatment with TT1_Ct_ decreased infarct volume to a similar extent, respectively 42% and 45% in female and male mice, differences between TMyc and TT1_Ct_-treated animals being statistically significant for both genders (\**p*<0.05 for female, \*\**p*<0.01 for male; Figure 7C). Interestingly, we observed a sex-dependent difference in neurological damage due to the ischemic insult. In male animals, changes in balance and motor coordination established in the beam walking test correlated with brain damage (Figure 7D). Thus, TMyc-treated mice presented a significant higher number of slips (10 ± 1) compared to TT1_Ct_-treatment (5 ± 1, \*\*\**p* < 0.001, n = 12), representing a 49% recovery. However, TMyc-treated female mice showed more discreet motor deficits than males (5 ± 1 versus 10 ± 1, \*\**p* < 0.01, n = 10), hindering the observation of a TT1_Ct_ effect with this test and suggesting a sexual dimorphism in the neurological outcome after photothrombotic ischemia.

**Figure 7.**
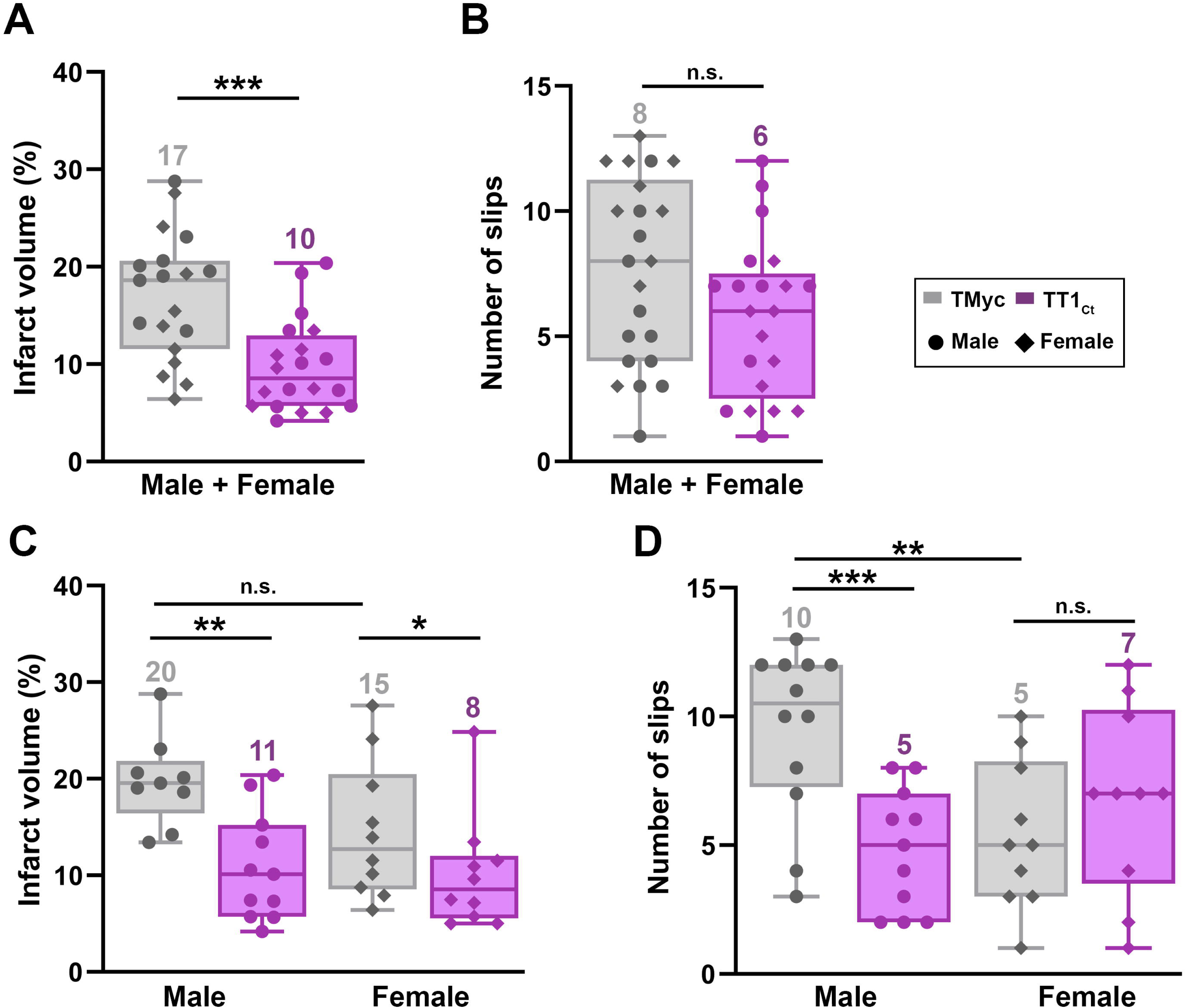
Treatment with TT1_Ct_ reduces infarct volume and motor coordination deficits in animals exposed to permanent ischemia. (A and C) Infarct volume of animals injected with TMyc or TT1_Ct_ (3 nmol/g) and sacrificed 24 h after damage induction, expressed as a percentage of the hemisphere volume. Individual data and box and whisker plots show interquartile range, median, minimum and maximum values. The mean value for each experimental group is also provided as a number on top of the corresponding plot. Results are given for the whole population (A) or disaggregated according to gender (C). Differences were analyzed by Student’s *t*-test (n = 20 for A, n = 9-10 for C). (B and D). Evaluation of balance and motor coordination. Number of contralateral hind paw slips were measured in male and female animals. As above, results are presented for the whole population (B) or disaggregated according to gender (D). Differences were analyzed by Student’s *t*-test (n = 21-22 for B, n = 10-12 for D).

In conclusion, these results demonstrate that peptide TT1_Ct_ is neuroprotective *in vivo* both in male and female mice subjected to permanent brain ischemia, reducing the infarct size in parallel to a decreased neuroinflammatory response. Furthermore, administration of TT1_Ct_ improves the neurological deficits resulting from ischemia in the male subpopulation. Altogether, these data support the importance of protein interactions established by the C-ter of neurotrophin receptor TrkB-T1 for reactive gliosis after brain damage, a process that has an impact on neurotrophic signaling and neuronal survival.

## Discussion

Glial cells reactivity and subsequent inflammation are common processes to ischemic stroke (Xu et al., 2020) and other acute or chronic neurodegenerative conditions (Paolicelli et al., 2022; Verkhratsky et al., 2023). These cells contribute to the pathophysiology of neurological diseases by protective and detrimental effects which imply complex and heterogeneous changes in glial cell morphology, gene expression and function. Consequently, the design of neuroprotective therapies targeted to glial cells, able to modulate neuroinflammation, is a priority but also a difficult task. Truncated neurotrophin receptor TrkB-T1, expressed both in neurons and glial cells, plays a relevant role in neuronal excitotoxicity (Gomes et al., 2012; Vidaurre et al., 2012) while, in astrocytes, promotes injury and edema formation after stroke (Colombo et al., 2024). Therefore, we hypothesized that TrkB-T1 function might be important for reactive gliosis and neurotoxicity, particularly through interactions with still undefined proteins established by the isoform-specific sequence. Accordingly, the modification of such interactions might prevent those pathological processes and, consequently, decrease the ischemic damage. In this work, we have designed peptides containing the TrkB-T1 C-ter sequence which modulate reactive gliosis and are strongly neuroprotective against *in vitro* and *in vivo* excitotoxicity. These Tat-derived peptides are able to cross the plasma membrane and, *in vivo*, the BBB, reaching the cortical region of both undamaged or damaged animals, where they are detectable for at least 4 h after i.v. injection.

One important mechanism of TT1_Ct_ action in excitotoxicity is maintenance of promoter activities dependent on CREB or MEF2, TFs involved in neuroinflammation due to brain injury (Wu et al., 2023; Yamaguchi et al., 2020; Yi et al., 2007) and key to neuronal survival/death (Hardingham et al., 2002; Lisek et al., 2023). Compared to neurons, the role played by CREB in astrocytes is not completely defined (Kim and Kaang, 2022) although it seems to regulate different adaptative transcriptional programs in both cell types (Pardo et al., 2017). Interestingly, expression of a constitutively active CREB form in astrocytes is neuroprotective in traumatic brain injury (Pardo et al., 2016). In our cell model of excitotoxicity, maintenance of CREB/MEF2 promoter activities by TT1_Ct_ action provokes relevant changes in levels of CREB and/or MEF2-regulated mRNAs encoding proteins important for astrocyte function. Thus, relative to cultures incubated with the control peptide and subjected to excitotoxicity, TT1_Ct_ induces a strong increase of CREB/MEF2-regulated *Bdnf* transcription (Lyons et al., 2012; Shieh et al., 1998; Tao et al., 1998) while it inhibits TrkB-T1 and GFAP expression. In cortical cultures, concurrent to decreased promoter activity, excitotoxicity reverts the normal pattern of *Ntrk2* alternative splicing, and favoursTrkB-T1 mRNA relative to TrkB-FL mRNA (Vidaurre et al., 2012). Treatment with TT1_Ct_, parallel to maintenance of CREB/MEF2 promoter activities, prevents TrkB-T1 mRNA increase induced by NMDA while it has no effect on TrkB-FL mRNA. *Gfap* transcription also seems to be negatively regulated by these TFs. Although, in our cortical cultures, we do not reproduce *Gfap* transcriptional activation induced by *in vivo* excitotoxicity, probably due to Ara C blockade of astrocyte proliferation, TT1_Ct_ significantly reduces this mRNA. An inverse association of CREB or MEF2D activation with GFAP expression has been described before in models of Alzheimer disease (Pugazhenthi et al., 2011) or isoflurane neuroprotection in ischemia/reperfusion (Zhang et al., 2021). These results are relevant because GFAP expression levels, alongside astrocyte reactivity, are generally considered as proportional to injury severity (Escartin et al., 2019).

We observed similar TT1_Ct_ effects in a preclinical model of stroke, where the process of excitotoxicity takes place *in vivo*. Very early after the ischemic damage (5 h), levels of TrkB-T1 and GFAP proteins increase in some cells of the infarcted area, particularly in the interface between ischemic and non-ischemic tissue which corresponds to developing glial scar. At later times (24 h), GFAP expression further increases in this area in TMyc-treated animals, mostly in cells also expressing C3, a marker of astrocytes with an inflammatory profile. In contrast, TT1_Ct_ strongly decreases GFAP+/C3+ cells both at 5 and 24 h of ischemic damage. Remarkably, TT1_Ct_ also downregulates macrophage BBB crossing into the injured tissue, in parallel to reduced IgG infiltration indicative of reduced BBB damage, as well as microglial Iba1 upregulation which takes place in the infarct core and the interface region between the ischemic and non-ischemic tissue. Globally, the results obtained *in vitro* and *in vivo* demonstrate the importance of TrkB-T1 signaling mediated by its C-ter sequence in modulation of the acute inflammatory response after stroke. Protein complexes formed by unprocessed TrkB-T1 or RIP TrkB-T1-ICD fragment with still unknown proteins might regulate the expression of proteins central to ischemia pathogenesis such as the truncated receptor itself, GFAP, C3 or Iba1. We do not presently know if this regulation only depends on CREB/MEF2 activities or additional proteins are involved. One possibility is that, similarly to Notch ICD, also a RIP product, TrkB-T1-ICD directly interacts with CREB and blocks this TF activity and expression of CREB-dependent genes (Hallaq et al., 2015). Anyway, by interfering TrkB-T1 interactions, peptide TT1_Ct_ might prevent these transcriptional changes, resulting in decreased microglia and astrocyte reactivity after an ischemic damage.

To reveal in a more integral way the protein interactions established by TrkB-T1 C-ter in neural cells and the relevance such proteins have in receptor signaling and function, we performed proteomic analysis with peptide Bio-sTT1_Ct_. It was important to determine the interactome established by TrkB-T1 C-ter inside cells not only in pathological but also non-pathological conditions. Except for RhoGDI, established as a TrkB-T1-interacting protein by affinity chromatography of astrocyte extracts with the isoform-specific 11 aa peptide (Ohira et al., 2005), no other TrkB-T1-interacting proteins had been identified before. This led to the notion that this short TrkB-T1 intracellular sequence might be unable to form stable protein interactions (Tessarollo and Yanpallewar, 2022). However, results presented here demonstrate that Bio-sTT1_Ct_ is able to establish a specific protein-interacting profile inside neural cells which, additionally, is susceptible to physiological and pathological conditions. A differential analysis of Bio-sTT1_Ct_-interactions in the three experimental conditions shows that, independently of treatment, this peptide interacts with proteins highly involved in gene expression regulation, including multiple proteins involved in processes such as RNA modification or splicing. One of them is adenosine deaminase acting on RNA (ADAR1), RNA editing enzyme central to regulation of short-term brain inflammation after ischemic injury (Cai et al., 2023). In stroke, ADAR1 is highly induced in astrocytes where it promotes proliferation and inflammatory cytokine production. Regarding BDNF treatment, we observe an enrichment of pathways mainly related with vesicle transport and mTOR regulation. It is particularly interesting the significant increase induced by BDNF in Bio-sTT1_Ct_-binding to neuron-enriched cholesterol 24-hydroxylase (CYP46A1), an enzyme that controls cholesterol turnover by converting it into 24S-hydroxycholesterol (24S-HC), a product that contributes to ischemic damage by potentiation of NMDAR activity (Sun et al., 2024). Finally, induction of excitotoxicity also produces an enrichment in pathways related with vesicle transport and, more specifically, interrelated pathways involved in axon guidance, signaling by RhoGTPases and innate immune response, including L1CAM signaling. RhoGTPases and related proteins are important regulators linking the activation of surface receptors to the organization of actin and microtubule cytoskeletons. They coordinate highly relevant cellular processes, including cell migration and polarity, cell cycle progression, gene transcription and neuronal survival and death (Stankiewicz and Linseman, 2014). They are also key in development of the ischemic damage, especially the morphological changes associated with reactive gliosis. Thus, upregulation of RhoA/ROCK signaling by ischemia contributes to transcriptional changes related to nuclear factor kappa B (NF-κB) activation and expression of chemokines genes, conducting to functional regulation of microglial cells and astrocytes, and neuroinflammation (reviewed in Lu et al., 2023).

One protein involved in RhoGTPase signaling that differentially and significantly interacts with Bio-sTT1_Ct_ in excitotoxicity is p21-activated kinase 5 (PAK5). It belongs to a family of Ser/Thr kinases which are downstream effectors of RhoGTPases Rac1 and Cdc42. They participate in different processes by cytoskeleton remodeling, promotion of gene transcription, included that of inflammatory cells, and cell survival (Civiero and Greggio, 2018). Thus, PAK proteins promote neuronal survival downstream Rac activation, as opposed to Rho promotion of neuronal death, by activation of mitogen activated protein kinases (MAPKs), and inhibition and enhancement respectively of pro-apoptotic and anti-apoptotic members of the Bcl-2 family (Johnson and D’Mello, 2005; Stankiewicz and Linseman, 2014). Interestingly, it has been recently discovered a P2Y12R-Rac-PAK-F-actin pathway used by microglia to alleviate neuronal cells from cytotoxic protein aggregates and regulate delivery of healthy mitochondria to burdened neurons through tunneling nanotubes (TNTs) (Scheiblich et al., 2024). These are connections established by microglia with neurons both in physiological and pathological conditions, critical to maintain neuronal function and brain health. Another protein central to TNT formation is actin-related protein 2/3 (Arp2/3), a multiprotein complex that modulates actin polymerization and depolymerization downstream of Rac. It is notable that, in this work, we have identified two Arp2/3 subunits (Actr2/Arp2 and Arpc3/p21-Arc) as differentially interacting with Bio-sTT1_Ct_ in the excitotoxic conditions. Finally, it has been demonstrated that, in response to spinal cord injury or cerebral ischemia, PAK5 synthesis and signaling is activated in axons, and these changes protect neurons from an axonal energy crisis. By reprogramming mitochondrial trafficking and anchoring, promoting replacement of damaged mitochondria with healthy ones, PAK5 facilitates neuronal survival and regeneration (Huang et al., 2021).

Other RhoGTPases-related proteins which differently increase their interaction with Bio-sTT1_Ct_ in excitotoxicity are the putative receptor of complement component 1 Q (C1qbp), C1q being secreted by reactive microglia and involved in the induction of neurotoxic reactive astrocytes (Liddelow et al., 2017), two proteins (Nup98 and Ranbp2) important for maintenance and/or assembly of the nuclear pore complex and nucleocytoplasmic transport, whose impairment is a hallmark of neurodegeneration (Ferreira, 2023), and engulfment and cell motility 2 (Elmo2) protein, an upstream Rac1 regulator required for phagocytosis and cell migration (Gumienny et al., 2001). Altogether, our results strongly suggest that overactivation of the NMDARs might alter the interactions established by TrkB-T1 C-ter with proteins involved in signaling by RhoGTPases, critical to cell morphology, neuronal survival and reactive gliosis. While some TrkB-T1-interaction are inhibited in excitotoxic conditions, others are strongly promoted. Displacement by Bio-sTT1_Ct_ of the latter could interfere signaling pathways associated to neuroinflammation and neurotoxicity, thus leading to neuroprotection.

In addition to RhoGTPase-related proteins, other proteins differentially interacting with Bio-sTT1_Ct_ in excitotoxicity might play important roles in inflammation and excitotoxicity. For example, calpain 5 (*Capn5*), whose expression is induced in reactive astrocytes in a model of ischemia/reperfusion and might enhance inflammation by caspase-11 activation (Chukai et al., 2024). We also find an increase in excitotoxicity of Bio-sTT1_Ct_-interaction with Septin-7, a cytoskeletal protein with GTPase activity involved in glioma progression (Hou et al., 2016). Interestingly, a proteomic study of human brain after stroke demonstrated specific changes in Septin-7 and some other proteins possibly related to TrkB-T1 signaling: Arp2/3, RhoGDI or complement protein C3 (Cuadrado et al., 2010), supporting the neuroprotective and anti-inflammatory potential of peptide TT1_Ct_.

The complexity of TrkB-T1 interactome allows us to propose that a combination of mechanisms affecting neurons and astrocytes alike could be responsible for TT1_Ct_ neuroprotective actions, demonstrated in excitotoxicity induced *in vitro* and an *in vivo* model of ischemia. In cortical cultures, this peptide is strongly neuroprotective when used at moderate concentrations (15 µM) and added before or at the time of the excitotoxic stimulus. This result is important from a preclinical and clinical perspective because, differently from cultured cells, excitotoxicity is not synchronous *in vivo*. By the time treatment is initiated in the ischemia model, penumbra neurons, our therapeutic target, would be exposed, if not properly protected, to secondary damage associated to infarct expansion. In fact, treatment with TT1_Ct_ 1 h after ischemic injury led to a significant reduction of the infarct volume both in male and female mice. Interestingly, we find sexual dimorphism in the post-ischemia motor coordination and balance impairments. Males show more pronounced motor deficits, which can be improved by TT1_Ct_-treatment, while females are less compromised despite having similar infarct volumes. Sexual dimorphism has been observed before in other studies evaluating motor activity and balance after CNS injury (Abbasian et al., 2022; Farooque et al., 2006), which has been related to a neuroprotective estrogen effect or a decreased neuroinflammatory response in female mice (Li et al., 2022). Therefore, future studies on the pathophysiology of ischemic brain damage and testing of therapeutic compounds should take into account the impact of biological sex.

In summary, we have designed a brain-accessible TrkB-T1-derived CPP which has proven to be relevant for the identification of the TrkB-T1 interactome in different biological conditions, and has multimodal effects affecting neurons and brain inflammation, as demonstrated *in vitro* and *in vivo*. After ischemic damage, this CPP induces neuroprotection and improves the neurological outcome, which might be relevant for the design of novel therapies for human acute stroke and other neurological conditions associated with excitotoxicity and neuroinflammation.

## Acknowledgements

We acknowledge funding from Agencia Estatal de Investigación (PID2019-105784RB-100/AEI/10.13039/501100011033/FEDER, UE and PID2022-137710OB-I00/AEI/10.13039/501100011033/FEDER, UE). The cost of publication has been paid in part by FEDER funds. A contract was funded associated to project PID2019-105784RB-100 (G.M.E-O) and L.U-T. was recipient of fellowships from Spanish Ministry of Universities “Ayuda del Programa de Formación de Profesorado Universitario” with code FPU22/01248 (2024-2028) and “Ayuda para el Fomento de la Investigación en Estudios de Master-UAM” fellowship from Universidad Autónoma de Madrid (2021-2022). We thank Conexión de Namomedicine CSIC for their support. We are also grateful to Drs. Gutiérrez (IdiPaz, HU La Paz), Lorrio (UAM), Sobrado (HU La Princesa), Avendaño and Negredo (Departamento de Anatomía, Histología y Neurociencia, UAM, Spain) for technical advice with the ischemia model. We acknowledge the contribution of Genomics and Microscopy Core Units (IIBM, CSIC) as well as the Proteomics Core Facility at Centro Nacional de Biotecnología (CNB, CSIC). Finally, we also thank Dr. M.C. Serrano (Instituto de Ciencia de Materiales de Madrid, CSIC) and members of our group for helpful discussions.

## Author Contribution

Conceptualization, M.D-G., L.U-T., G.M.E-O. and G.S.T.; Methodology, M.D-G., L.U-T., G.S.T. and G.M.E-O.; Formal Analysis, M.D-G., G.S.T., G.M.E-O. and L.U-T.; Investigation, L.U-T., G.S.T. and G.M.E-O.; Writing – Original Draft, M.D-G. and L.U-T.; Writing – Review & Editing, M.D-G., L.U-T. and G.S.T.; Visualization, M.D-G. and L.U-T.; Supervision, M.D-G.; Funding Acquisition, M.D-G.

## Material and Methods

All reagents, antibodies, culture products, recombinant DNAs and primers are described in the Key Resources Table.

### Experimental models

Animal procedures were performed following European Union Directive 2010/63/ EU and were approved by the CSIC and Comunidad de Madrid (Ref PROEX 276.6/20) ethics committees. Housing facilities were approved by Comunidad de Madrid (#ES 280790000188) and conform to official regulations. Animals had standard health and immune status and were looked after by professional caretakers daily. Male and female mice were kept in groups of up to five in standard IVC cages while 1–2 pregnant rats occupied standard cages, always containing bedding and nesting materials. Animals were under controlled lighting conditions (12 h light cycles), relative humidity and temperature. Irradiated food and water were provided *ad libitum*. We tried to minimize animal suffering and reduce the number of sacrificed animals.

### Mice model of ischemia by photothrombosis to study *in vivo* excitotoxicity

Permanent focal ischemia was induced in the cerebral cortex of adult male and female Balb/cOlaHsd mice (23–31 g; 10–16 weeks of age; Envigo Rms, Spain S.L.) by microvascular photothrombosis. This model mimics embolic or thrombotic occlusion of small arteries, which is frequently found in human stroke, and causes a focal brain damage with histological and MRI correlations to human patterns (Pevsner et al., 2001). Animals were allowed to acclimatize to our facilities for at least 1 week after delivery, before ischemic induction, and were kept with the same cage mates, and daily examined to check their health status. Mice were anesthetized with isoflurane (5% for induction, 2% for maintenance in oxygen; Abbot Laboratories, Madrid, Spain) and then placed in a stereotaxic frame (Narishige Group, Tokyo, Japan). Body temperature was maintained at 36–37°C using a self-regulating heating blanket (Cibertec, Madrid, Spain). A midline scalp incision was made, the skull was exposed, and both Bregma and Lambda points were identified. A cold-light (Schott KL 2500 LCD; Schott Glass, Mainz, Germany) with a fiber optic bundle of 1.5 mm in diameter was centered using a micromanipulator on the right side, at 0.2 mm anterior and 2.0 mm lateral (+0.2 AP, +2 ML) relative to Bregma. Afterward, the photosensitive dye Rose Bengal (7.5 mg/ml, prepared in sterile saline; Sigma-Aldrich) was administered by retro-orbital injection of the venous sinus, for intravenous (i.v.) vascular access, to a body dose of 20 mg/kg. Five minutes later, the brain was illuminated (600 lms, 3,000 K) through the intact skull for 10 min. Brain injury encompasses damage to vascular endothelium, platelet activation, and subsequent microvascular thrombotic occlusion of the irradiated region (Watson et al., 1985). According to the Paxinos mouse brain atlas, the areas underneath this stereotaxic position that result irradiated are the primary motor cortex and the primary somatosensory cortex (hindlimb and forelimb). Once completed surgery, the incision was sutured, and mice were allowed to recover.

For neuroprotection experiments, a single dose (3 nmol/g) of peptides TMyc or TT1_Ct_ (> 95% purity; GenScript), solubilized as 2.5 mM solutions in 0.9% NaCl, we retro-orbitally injected 1 h after damage initiation. In order to improve their plasma stability, these peptides are N-and C-ter modified by, respectively, acetyl and amide groups. Mice were not subjected to other procedures before ischemia and were naive to drug or peptide treatment. Animals were randomly allocated to the experimental groups and the researchers doing the experiments were blind respect to treatment. There was not a previous estimation of sample size and experiments were independently carried out as indicated. Twenty-four hours after damage induction, animals completed the beam walking test and were subsequently sacrificed by cervical dislocation to measure the infarct volume, as indicated below. Before that, brains were sectioned into serial 1-mm-thick coronal slices using a mouse brain matrix (Stoelting, Wood Dale, IL, USA), and sections were completely stained with 2% TTC at room temperature to visualize the cortical infarcts. For immunohistochemistry, animals were deeply anesthetized 5 or 24 h after brain damage and intracardially perfused with cold PBS and 4% paraformaldehyde in PBS as indicated below.

### Primary cultures of rat cortical neurons to study *in vitro* excitotoxicity

Primary neuronal cultures were prepared from cerebral cortex of 18-day-old Wistar rat embryos (E18), both genders being indistinctly used. The dissected cerebral cortices were mechanically dissociated in culture medium (Minimum Essential Medium, MEM, Life Technologies) followed by seeding of the cell suspension at a density of 1.1×10^6^ cells/ml, prepared in the same medium supplemented with 22.2 mM glucose, 0.1 mM glutamax, 5% fetal bovine serum (FBS), 5% donor horse serum (HS), 100 U/ml penicillin and 100 µg/ml streptomycin. Before seeding, the plates were treated overnight at 37°C with 100 µg/ml poly-L-lysine and 4 µg/ml laminin. The cells were incubated at 37°C in an atmosphere of 5% CO_2_ and 95% humidity. Unless otherwise indicated, glial growth was inhibited after 7 days *in vitro* (DIVs) by adding 10 µM 5 cytosine β-D-arabinofuranoside (AraC) and experimental treatments took place after 12 DIVs. These cultures present a high percentage of neurons, combined with glial cells, mostly astrocytes (Vidaurre et al., 2012).

### Measurement of infarct volume

Sections stained with 2% TTC, as before indicated, were subsequently scanned by both rostral and caudal sides. These images were analyzed using ImageJ software by an observer blinded to experimental groups. After image calibration, delineated areas of ipsilateral and contralateral hemispheres, and the infarcted region (unstained area) were measured. Considering slices thickness, the corresponding volumes were calculated and corrected for edema’s effect, estimated by comparing total volumes of hemispheres. The corrected infarct volumes were expressed as percentage relative to the contralateral hemisphere, to correct for normal size differences between different animals. For each animal, the mean of results obtained for rostral and caudal sides was calculated.

### Beam walking test

Motor coordination and balance were evaluated in mice right before and 24 h after the ischemic insult by measuring the number of contralateral hind paw slips in the beam walk apparatus. Mice had to walk through a narrow beam (1 m × 1 cm × 1 cm) placed 50 cm above the tabletop, going from an aversive stimulus (60-W light bulb) to a black goal box with nesting material. Slips taking place in a previously selected central beam segment (50 cm long) were counted. Before damage induction, mice were allowed to cross the beam once, to get acquainted with the test, which they repeated 24 h after photothrombosis, immediately before sacrifice.

### Induction of neuronal excitotoxicity

To induce excitotoxicity, cultures were incubated with NMDA (100 µM) and its co-agonist glycine (10 µM), a treatment herein denoted simply as NMDA. These NMDAR coagonists induce a strong excitotoxic response in the mature neurons present in the culture but have no effect on astrocyte viability (Choi, 1985; Choi et al., 1987). When indicated, primary cultures were preincubated for 30 min with the indicated concentrations of Tat-derived CPPs before NMDA treatment, the peptides being kept in the medium along treatment.

### Assessment of neuronal injury in cortical cultures

Thiazolyl blue formazan (MTT) reduction assay was used to determine cell viability. At the end of the different treatments, 0.5 mg/ml MTT was added to the medium, and after 2 h of incubation at 37°C the formazan salts formed were solubilized using DMSO. The results were quantified spectrophotometrically at 570 nm. As primary cortical cultures are formed by neurons and glial cells, we determined the magnitude of glia viability on the total values by applying 400 µM NMDA and 10 µM glycine to sister cultures 24 h before MTT assay. These conditions induce a complete neuronal death but no glial damage. Thus, after subtracting this absorbance value, the results only represent the viability of neurons present in cultures. Each independent experiments included technical triplicates for every treatment condition. In addition, a minimum of 4 completely independent experiments were performed and analyzed as indicated in figure legends. When indicated, the provided viability data correspond to total cell values (neurons and glial cells.

### Synthetic peptides

Synthetic cell-penetrating peptides (> 95% purity; GenScript) were used for treatment of primary cortical cultures and mice. All these CPPs contain a HIV-1 Tat sequence (11 aa) linked to a specific TrkB-T1 or c-Myc sequence as indicated (Key Resources Table). Three of these peptides (Bio-TMyc, Bio-sTT1_Ct_ and Bio-TT1Ct) contain a biotin molecule at the N-ter, which serves several purposes: peptide labelling, stabilization, and competence to isolate proteins interacting with TrkB-T1 taking place inside cultured cells. In the other two peptides (TMyc and TT1Ct), the N-ter is acetylated while all five CPPs have their C-ter amidated. The concentration of use is indicated for each experiment.

### Pull down assays

Cell cultures were lysed at 4°C in NP-40 lysis buffer (1% NP-40, 20 mM Tris HCl pH 8.0, 80 mM NaCl, 20 mM EDTA) containing protease and phosphatase inhibitors (Complete protease and PhosSTOP phosphatases inhibitor cocktail tablets, Roche). Protein concentration was determined using BCA Protein Assay Kit (Thermo Fisher). Equal amounts of protein were combined with streptavidin-agarose beads previously washed with NP-40 lysis buffer and incubated at room temperature for 1.5 h with agitation. After sedimentation, beads were washed 3 times with the NP-40 lysis buffer to remove possible non-specifically bounded proteins. Next, peptides and their interacting proteins were released from beads by incubation at 50°C for 40 min in RIPA modified buffer (50 mM Tris HCl pH 8.0, 150 mM NaCl, 1% sodium deoxycholate, 1% NP-40) combined with protease and phosphatase inhibitors as before. After centrifugation, the supernatant containing the interacting proteins was collected and subjected to LC-ESI-MS/MS (HR, medium gradient) protein identification using a mass spectrometer (Orbitrap Exploris 240) and Proteome Discoverer software for analysis (CNB Proteomics Facility, CSIC).

### Immunoblot analysis

After washing with cold PBS, cultured cells were lysed in RIPA buffer (50 mM Tris-HCl pH 8.0, 150 mM NaCl, 1% sodium deoxycholate, 1% NP-40, 0.1% SDS, 1 mM DTT), supplemented with protease and phosphatase inhibitors as above, for 30min at 4°C. Then, the lysate was sonicated using a Bioruptor apparatus and centrifugated at 4°C for 20 min at 10,000 rpm. The protein concentration in lysates was established as before and total cell-lysates were denatured in SDS-sample buffer followed by heating at 95° for 5 min. Equal amounts of total lysates were resolved in Tris-Glycine SDS-PAGE and transferred to a nitrocellulose membrane (GE Healthcare) in 25 mM Tris HCl pH 8.3, 250 mM glycine and 10% methanol, using an electric current of 400 mA for 70 min. After transfer, membranes were stained with a Ponceau S solution to check for its efficacy. Then, membranes were blocked for 30 min with a 5% non-fat dry milk solution in Tris Buffered Saline-Tween (TBS-T, 20 mM Tris HCl pH 7.5, 137 mM NaCl, 0.05% Tween-20) and incubated overnight at 4°C with primary antibodies. Next, membranes were washed with TBS-T and incubated with the appropriate anti-rabbit or anti-mouse peroxidase-conjugated secondary antibodies (Bethyl) for 1 h. To conclude, immunoreactivity was detected using Clarity Western ECL Blotting Substrate (BioRad) and band intensity was quantified by densitometric analysis (Photoshop, Adobe). Levels of the protein of interest were normalized using those of neuron-specific enolase (NSE) present in the same sample and expressed relative to values obtained in their respective controls, arbitrarily given a 100% value. NSE was used as a neuronal loading control since it is not affected by NMDA treatment. Multiple independent experiments were carried out and quantitated as detailed in the figure legends.

### Cell transfection and gene reporter assays

The plasmids used in these experiments contained minimal CREB or MEF2 response elements upstream the firefly luciferase reporter gene (respectively, pCRE or pMEF2; see details in Key Resources Table). Primary cortical cultures, without the AraC treatment, were transfected at 11 DIVs with 0.4 µg of plasmids using Lipofectamine 2000 (Life Technology). DNA-liposomes complexes were prepared according to the manufacturer instructions in neurobasal medium (Thermo Fisher Scientific) with 1 mM glutamax (Gibco, Thermo Fisher Scientific) and added to the cell culture. After 2 h of transfection, this mix was replaced with conditioned medium collected and saved at the beginning of the experiment, and incubation proceeded to complete 24 h of transfection. Finally, protein extracts were obtained using *Passive Lysis Buffer* (Promega, Cat# E1941) and luciferase activity was detected by a luminometer (GloMax 96 microplate luminometer, Promega), using REAP buffer (25 mM glycylglycine, 15 mM SO_4_Mg, 4 mM EGTA, 15 mM potassium phosphate pH 7.8, 3.3 mM ATP, 1 mM DTT, 75 µM luciferin). For quantitation, 5-7 totally independent experiments, each one including technical quadruplicates for every condition, were repeated as indicated.

### RNA extraction and qPCR assay

Total RNA was extracted using QIAcube technology and treated with DNases before cDNA synthesis by the “High Capacity cDNA Reverse Transcription Kit” (Applied Biosystems). A 7900HT Fast real-time PCR system (Applied Biosystems) was used for SYBR green gene expression assays, with the primers indicated in Table 1. PCR conditions were 10 min at 95°C, followed by 40 cycles of 15 s at 95°C and 60 s at 60°C. For each independent experiment, we made a specific standard curve for every gene and technical triplicates were prepared for every sample. Then, we performed a total of 5 completely independent experiments and, for each of them, data were normalized with housekeeping genes, neuronal specific enolase (NSE, for genes having neuronal expression) or glyceraldehyde-3-phosphate dehydrogenase (GAPDH, for GFAP, only expressed in glial cells).

### Peptide visualization in primary cultures and immunocytochemistry

Primary cultures, seeded at half the concentration previously indicated, were grown for 13 DIVs on coverslips previously treated overnight at 37°C with poly-L-lysine and laminin as before. Next, cells were incubated for 1 h with biotin-conjugated peptides Bio-TMyc or Bio-sTT1_Ct_ (25 µM) or left untreated, and then fixed for 30 min with 4% paraformaldehyde in PBS. The fixed cells were washed several times with PBS to eliminate the remaining paraformaldehyde before they were blocked and permeabilized for 4 h with 4% goat serum and 0.5% Triton X-100 prepared in PBS. Afterwards coverslips were incubated overnight at 4°C with the corresponding primary antibodies, diluted in the blocking solution. After washing with PBS, the coverslips were incubated with the corresponding secondary antibodies for 1 h at room temperature and then 10 min with DAPI, to stain DNA. Finally, the samples were mounted in Prolong Diamond. The images were acquired using an inverted Zeiss LSM 710 laser confocal microscope (Jena, Germany) with a 40x or 63x Plan-Apochromatic oil immersion objective. Images correspond to single sections and were normalized for each color separately and processed for presentation with ImageJ (NIH Image) and Fiji.

### Peptide visualization and immunohistochemistry in brain cortex

Vehicle (saline), Bio-TMyc or Bio-TT1_Ct_ (3 nmol/g) were retro-orbitally injected in mice one hour after the induction of photothrombotic damage or, alternatively, undamaged animals. In the first case, animals were sacrificed after completing 5 h of ischemic damage while, for the second group, the sacrifice took place 30 min after peptide administration. In all cases, animals were deeply anesthetized and intracardially perfused with cold PBS and 4% paraformaldehyde in PBS. Brains were post-fixed in the same fixative at 4°C for 24 h and cryoprotected in 30% sucrose for 48 h at 4°C. Coronal frozen sections (30 µm thick) obtained using a cryostat (Leica, Heidelberg, Germany) were incubated in blocking solution (10% goat serum, 0.5% Triton X-100 in PBS) for 3 h at room temperature, followed by Fluorescein Avidin D (200 µg/ml), and DAPI (5 µg/ml) prepared in 4% goat serum for 1 h. When indicated, sections were incubated for 3 h with anti-NeuN in 4% goat serum right after blocking and, after washing, with Alexa Fluor 647-conjugated antibodies together with Fluorescein Avidin D and DAPI as before indicated. Finally, sections were mounted and dried on slides, and cover slipped with Prolong Diamond. Image acquisition was performed using an inverted laser confocal microscope as before with a 63x or 40x objective Plan-Apochromatic oil immersion objective. Images were processed as described and correspond to single sections. Background was subtracted using vehicle-injected animals.

### Immunohistochemistry

The brains of animals injected one hour after the initiation of the photothrombotic damage with vehicle (saline), TMyc or TT1_Ct_ (3 nmol/g) were processed and cryoprotected as before indicated. The duration of damage before sacrifice was 5 h (male and female) or 24 h (male). Coronal frozen sections were incubated with blocking solution as described in flotation for 1 h at room temperature and then overnight with primary antibodies diluted in 4% goat serum and 0.3% Triton X-100 in PBS at 4°C. After washing, slides were incubated 1 h with Alexa Fluor 546-conjugated goat anti-rabbit or AlexaFluor 647-conjugated goat anti-mouse secondary antibodies to visualize the primary antibodies and DAPI (5 μg/ml). Only for Iba1 staining, blocking solution was 2.5% BSA, 5% goat serum, 0.5% Triton X-100 in PBS, and sections were incubated with primary antibody 24 h at RT followed by washing and Alexa Fluor 546-conjugated goat anti-rabbit and DAPI incubation for 1 h. Sections were then washed in PBS, mounted on slides, dried for 15 min at 37°C on a hot plate, or air dried overnight and then cover slipped with DPX or Prolong Diamond. Confocal images were acquired with a 40x Plan-Apochromatic oil immersion objective and processed for presentation as above described.

### Proteomic analysis

Proteomic analysis was performed in ProstaR (1.34.6) software tool (Wieczorek et al., 2017), using the protocol for protein-level analysis (Wieczorek et al., 2019). Proteomic abundances from 4 independent experiments were log-transformed and filtered following quality parameters: proteins present in at least >50% of one of the conditions and identified by at least 2 unique peptides. After, data was normalized by LOESS (locally estimated scatterplot smoothing, k=0.7), and POV (Partially observed values) and MEC (Missing in entire condition values) imputations were made using, respectively, KNN (k-nearest-neighbor k=10) and det.quantile (Quantile imputation using quantile=2,5). Differential analysis was performed using limma (3.16). Multiple comparison p-values where calibrated using slim algorithm and the FDR (False discovery rate) is indicated for each comparison. For PCA (Principal Component Analysis), filtered, normalized and imputed data was used corrected for batch effect using removeBatchEffect {limma} function. First and second principal components are presented. For volcano plot representation ggplot2 (3.5.1) software was used, and for heatmap presentation ComplexHeatmap (2.18.0) software. Enrichment analysis for GO (Gene Ontology terms) and Reactome (Reactome.db) pathway analysis the string.db (12.0) tool was used.

### Statistical analysis

Except for box and whiskers plots, data are expressed as mean ± standard error of the mean (SEM) of at least four independent experiments. The details of the number of completely independent experiments done (n) and the specific statistical test applied can be found in each respective figure legend. For viability, mRNA and gene reporter assays, technical replicates were included in each independent experiment. Treatment assignation was performed at random. Statistical analysis was performed in GraphPad Prism 8.0.2.The normality of the data was analyzed by Saphiro-Wilk test. In all cases *p*-value significance is considered as: \**p*<0.05, \*\**p*<0.01, \*\*\**p*<0.001, \*\*\*\**p*<0.0001. A *p*-value larger than 0.05 is considered non-significant (n.s.).

---

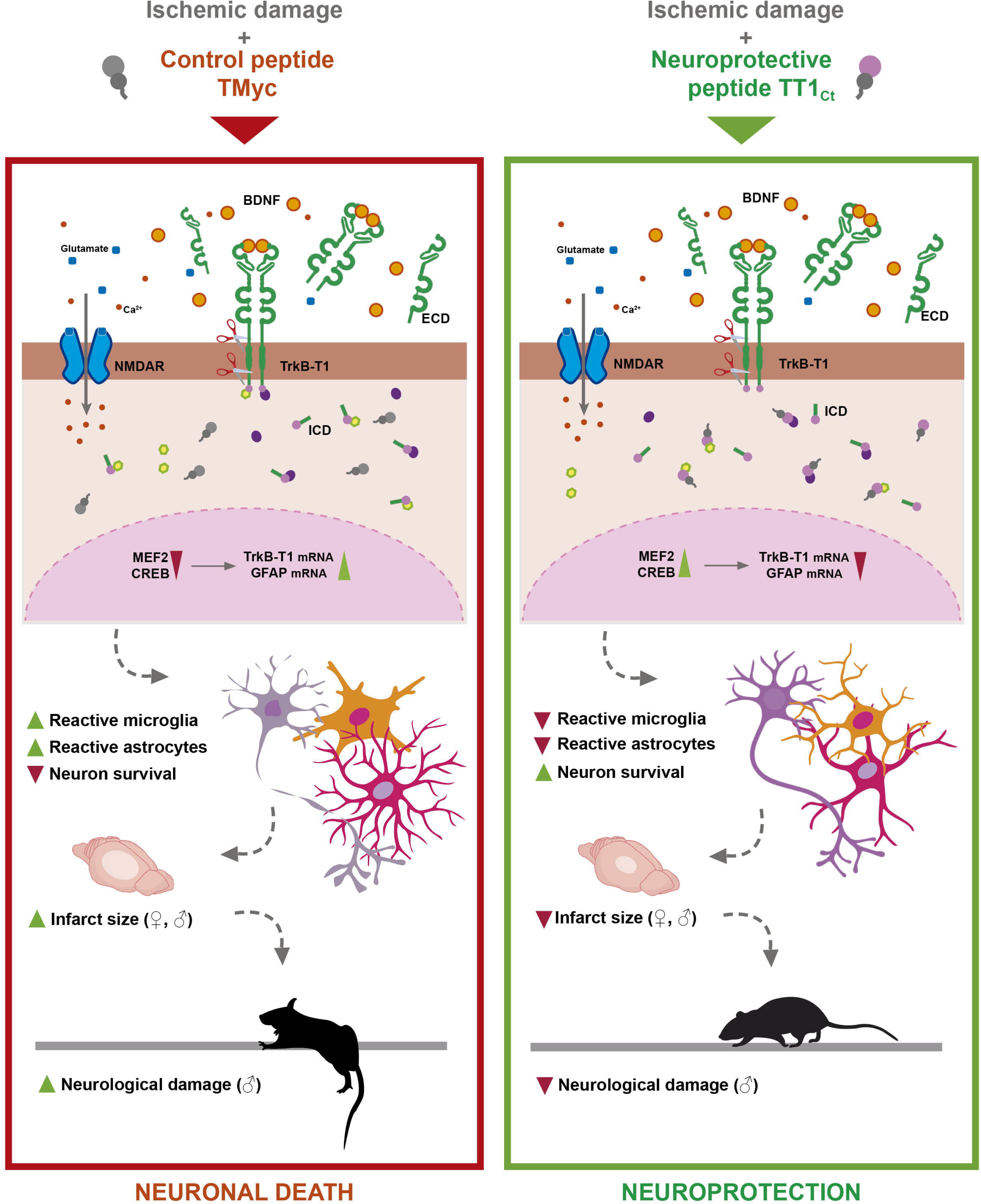

