## Supplementary Files for "Interactions established by isoform-specific TrkB-T1 sequences govern inflammatory response and neurotoxicity in stroke"

#### Supplementary Figures

**Figure S1.** Biological processes associated with Bio-sTT1$_{Ct}$ protein interactions.

**Figure S2.** Total RhoGDI1 levels are not affected by BDNF or NMDA treatment.

**Figure S3.** Neuroprotection due to TT1$_{Ct}$ preincubation is maintained when applied at the time of damage induction.

**Figure S4.** Bio-TT1$_{Ct}$ and Bio-TMyc can cross the BBB of undamaged animals.

**Figure S5.** Analysis of TrkB isoform expression in damaged brain of animals treated with Bio-TMyc.

**Figure S6.** Leakage of mouse immunoglobulins to the brain cortex due to BBB breakage is decreased in TT1$_{Ct}$-treated animals.

#### Supplementary Tables

**Table S1.** Selected Bio-sTT1$_{Ct}$-interacting proteins showing significantly altered levels after BDNF treatment compared to the basal conditions.

**Table S2.** Selected Bio-TT1$_{Ct}$-interacting proteins showing significantly increased levels after NMDA treatment compared to BDNF stimulation.

**Table S3.** Selected Bio-TT1$_{Ct}$-interacting proteins showing significantly altered levels after NMDA treatment compared to the basal conditions.

**Table S4.** *Arhgdia* (RhoGDI1) and group of selected Bio-TT1$_{Ct}$-interacting proteins included in the Reactome function “Signaling by RhoGTPases”.

**Key Resources Table**

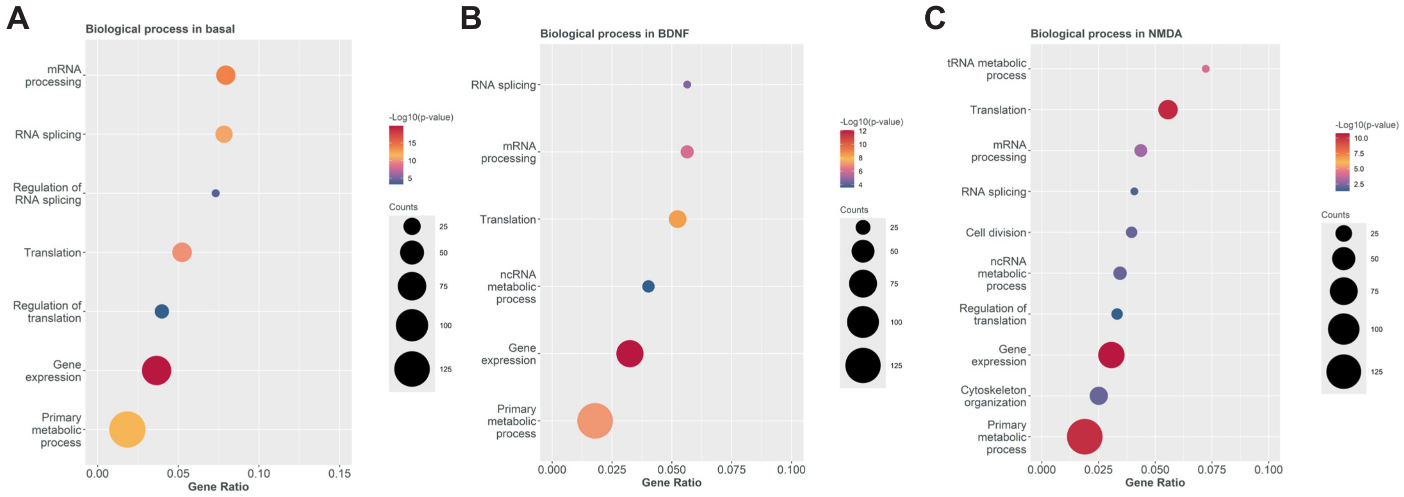


**Figure S1. Biological processes associated with Bio-sTT1$_{Ct}$ protein interactions.**
Representation of the most relevant Gene Ontology (GO) terms related to biological processes for selected proteins interacting with Bio-sTT1$_{Ct}$ in basal conditions **(A)** or after treatment with BDNF **(B)** or NMDA **(C)**. The dot size represents the number of proteins from our data set related to each process. Dots are colored according to their statistical significance, which is set by a color scale referring to -log10 (adjusted p-value).

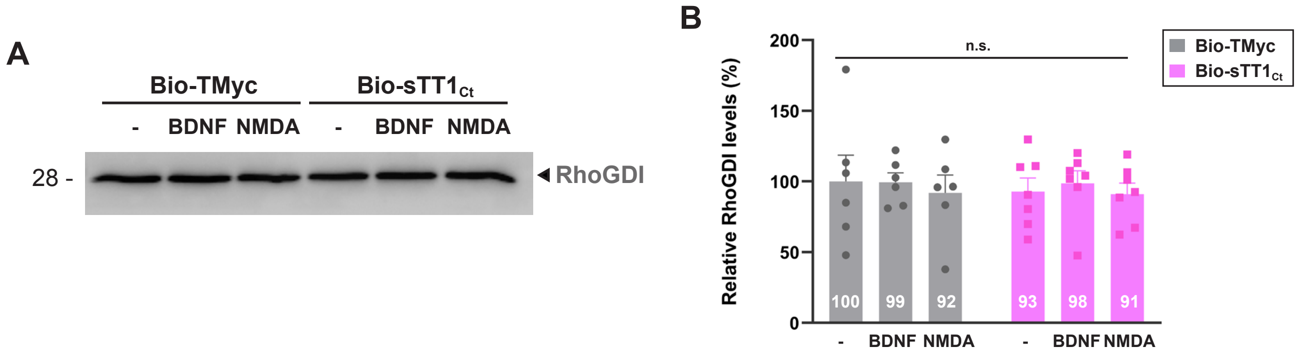


**Figure S2. Total RhoGDI1 levels are not affected by BDNF or NMDA treatment. (A)** Western blot analysis of cortical cultures incubated with Bio-sTT1$_{Ct}$ or Bio-TMyc (25 μM) for 30 min before treatment with BDNF (100 ng/mL) or NMDA for 30 min. A representative experiment is presented showing RhoGDI levels in total lysates. **(B)** Quantitation by densitometric analysis of RhoGDI total levels. Means ± SEM are presented relative to basal conditions with Bio-TMyc (100%). Data were analyzed by ANOVA test followed by *post-hoc* Tukey's test, n = 6-7.

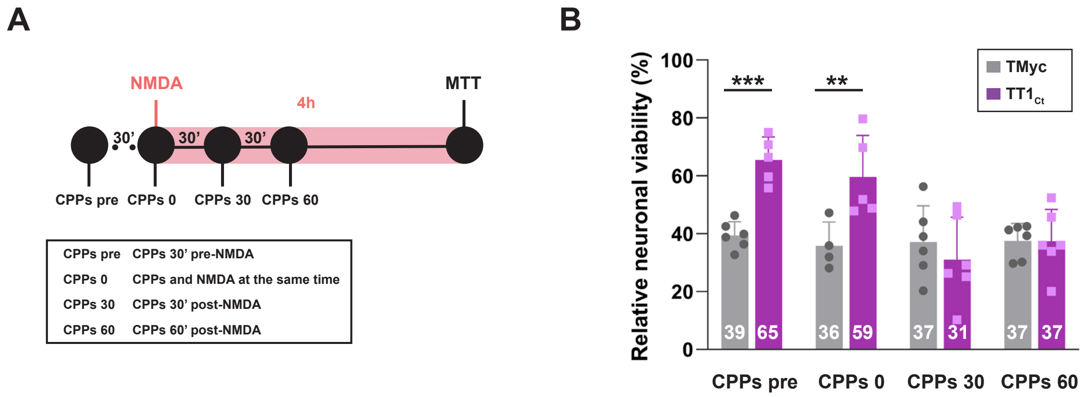


**Figure S3. Neuroprotection due to TT1$_{Ct}$ preincubation is maintained when applied at the time of damage induction. (A)** MTT assay design to evaluate TT1$_{Ct}$ neuroprotection in primary cortical cultures treated with CPPs at different time points of damage induction. TMyc or TT1$_{Ct}$ were added 30' before (CPPs pre), at the same time (CPPs 0), 30 min (CPPs 30) or 60 min (CPPs60) after treatment with NMDA (100 µM). Neuronal viability was established after 4 h of excitotoxicity. **(B)** Neuronal viability in cultures treated as above indicated. Individual results and means ± SEM are presented relative to the values obtained for the untreated cells (100%). Data were analyzed using two-way ANOVA test followed by *post hoc* Bonferroni test, n = 4-8.

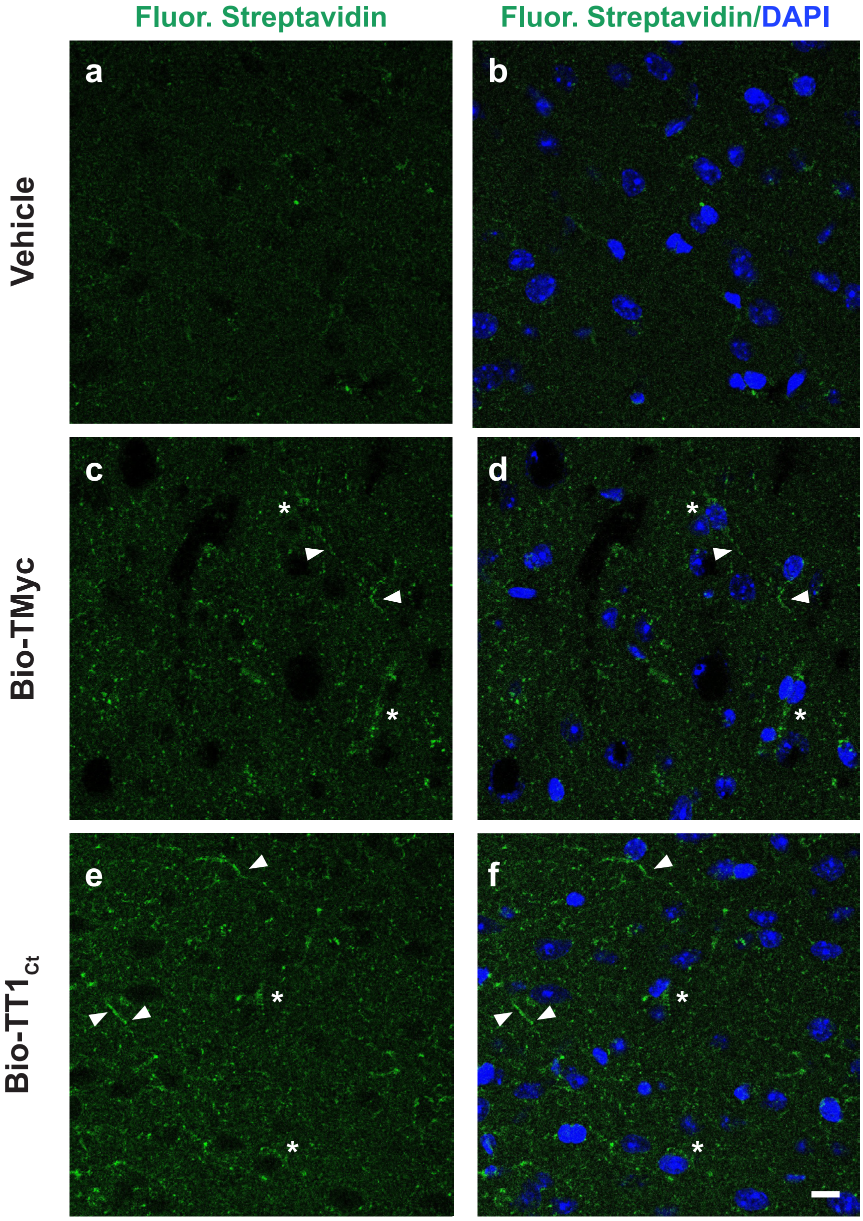


**Figure S4. Bio-TT1$_{Ct}$ and Bio-TMyc can cross the BBB of undamaged animals.** Analysis of biotinylated-peptide delivery to undamaged male mice cortex. Bio-TT1$_{Ct}$ and Bio-TMyc (3 nmol/g) were injected, and animals were sacrificed 30 min after. Detection was made in coronal sections by Fluorescein Avidin D (green). Peptide delivery was observed in cell projections (arrowheads) and bodies (asterisks) of cortical cells. Representative confocal microscopy images of cortical areas correspond to single sections. Scale bar, 20 µm.

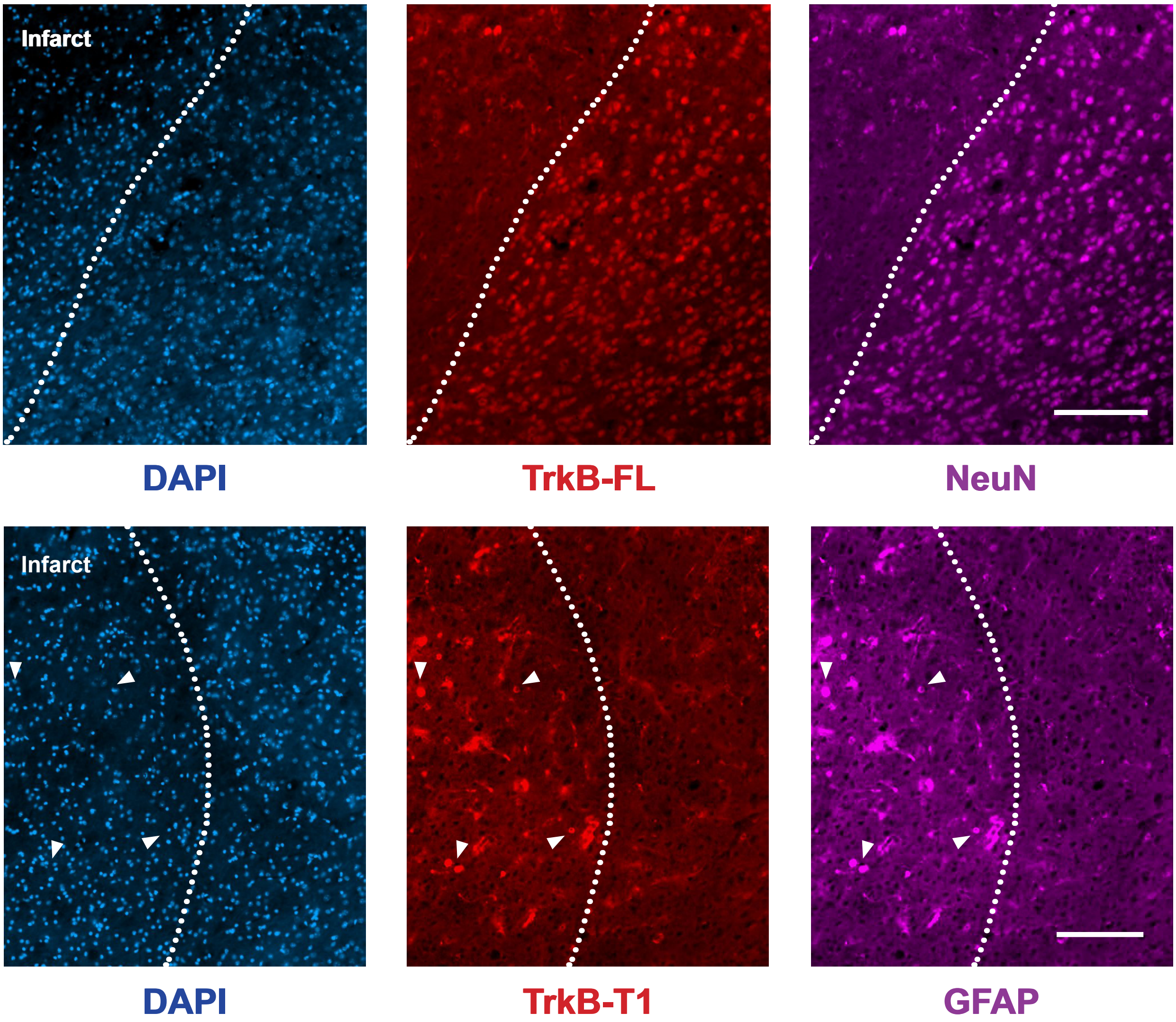


**Figure S5. Analysis of TrkB isoform expression in damaged brain of animals treated with Bio-TMyc.** Immunohistochemistry of brain coronal sections prepared from male animals i.v. injected with Bio-TMyc (3 nmol/g) 1 h after damage initiation and sacrificed after 5 h of injury. Sections were stained with isoform-specific TrkB antibodies (TrkB-FL and TrkB-T1), NeuN, GFAP and DAPI. Representative cell observer images of cortical areas of the infarct border are shown. Arrows indicate concurrent increased expression of TrkB-T1 and GFAP in astrocytes. Scale bar, 50 μm.

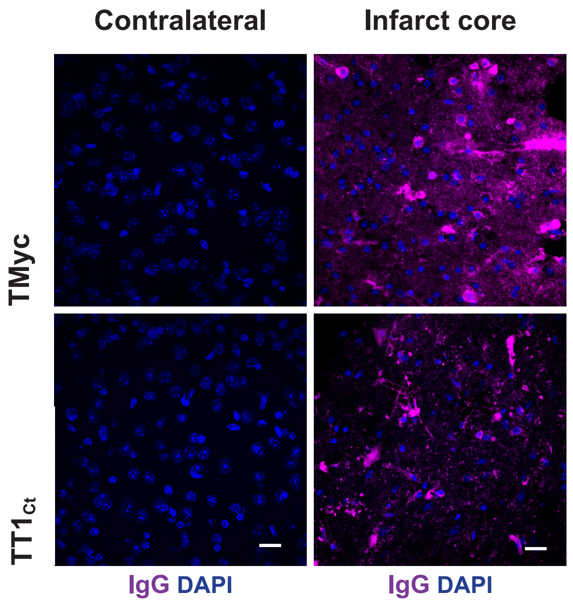


**Figure S6. Leakage of mouse immunoglobulins to the brain cortex due to BBB breakage is decreased in TT1$_{Ct}$-treated animals.** Brain coronal sections of male animals sacrificed 24 h after insult were analyzed by immunohistochemistry with an anti-mouse secondary antibody. Heavy staining of blood vessels, infiltrated cells and high backgrounds were observed in the ischemic brain of animals injected with the control peptide and decreased in TT1$_{Ct}$-treated animals. Representative images correspond to single sections. Scale bar, 20 µm.

**Table S1. Selected Bio-sTT1$_{ct}$-interacting proteins showing significantly altered levels after BDNF treatment compared to the basal conditions**

| Uniprot ID | Gene name | logFC BDNF vs Basal | p-value BDNF vs Basal | Description | Confidende | nº of unique peptides |
|---|---|---|---|---|---|---|
| A0A8I5Y1J2 | | 2.526 | 0.012 | Peptidyl-prolyl cis-trans isomerase OS=Rattus norvegicus OX=10116 GN=ENSRNOG00000067128 PE=3 SV=1 | High | 2 |
| A0A8I5Y9S3 | *Mrpl48* | -1.393 | 0.025 | Mitochondrial ribosomal protein L48 OS=Rattus norvegicus OX=10116 GN=Mrpl48 PE=1 SV=1 | High | 3 |
| A0A8I5ZLV1 | *Mtx2* | 2.616 | 0.031 | Metaxin 2 OS=Rattus norvegicus OX=10116 GN=Mtx2 PE=1 SV=1 | High | 2 |
| A0A8I5ZS37 | *Gak* | 1.724 | 0.028 | Cyclin G associated kinase OS=Rattus norvegicus OX=10116 GN=Gak PE=1 SV=1 | High | 3 |
| A0A8I5ZTG0 | *Cdc23* | -0.776 | 0.040 | CDC23 (Cell division cycle 23, yeast, homolog), isoform CRA_b OS=Rattus norvegicus OX=10116 GN=Cdc23 PE=1 SV=1 | High | 6 |
| A0A8I6A326 | *Rbm6* | -1.285 | 0.011 | RNA binding motif protein 6 OS=Rattus norvegicus OX=10116 GN=Rbm6 PE=1 SV=1 | High | 3 |
| A0A8I6A9H6 | *Rnps1* | 1.442 | 0.044 | RNA-binding protein with serine-rich domain 1 OS=Rattus norvegicus OX=10116 GN=Rnps1 PE=3 SV=1 | High | 3 |
| A0A8I6AKZ9 | *Actr1b* | 1.323 | 0.006 | Actin related protein 1B OS=Rattus norvegicus OX=10116 GN=Actr1b PE=1 SV=1 | High | 2 |
| A0A8I6AN44 | *Nfasc* | -1.092 | 0.027 | Neurofascin OS=Rattus norvegicus OX=10116 GN=Nfasc PE=3 SV=1 | High | 9 |
| A0A8I6GIS7 | *Relch* | -1.207 | 0.031 | RAB11 binding and LisH domain, coiled-coil and HEAT repeat containing OS=Rattus norvegicus OX=10116 GN=Relch PE=1 SV=1 | High | 2 |
| A0A8L2RBV8 | *Rpl39-ps13* | 0.864 | 0.038 | Ribosomal protein L39 OS=Rattus norvegicus OX=10116 GN=Rpl39 PE=1 SV=1 | High | 2 |
| F7EN52 | *Cyp46a1* | 2.113 | 0.027 | Cytochrome P450, family 46, subfamily a, polypeptide 1 OS=Rattus norvegicus OX=10116 GN=Cyp46a1 PE=3 SV=3 | High | 3 |
| P62893 | *Rpl39* | 0.864 | 0.038 | 60S ribosomal protein L39 OS=Rattus norvegicus OX=10116 GN=Rpl39 PE=1 SV=2 | High | 2 |
| Q5RJT2 | *Ftsj3* | 1.666 | 0.040 | pre-rRNA 2'-O-ribose RNA methyltransferase FTSJ3 OS=Rattus norvegicus OX=10116 GN=Ftsj3 PE=1 SV=1 | High | 3 |
| Q66H79 | *Trim32* | 1.694 | 0.044 | Tripartite motif protein 32 OS=Rattus norvegicus OX=10116 GN=Trim32 PE=1 SV=1 | High | 2 |

Uniprot ID and gene name are presented for each protein, together with its logFC obtained after comparison of BDNF vs basal conditions. Individual p-values for each protein are also shown. Protein description together with quality parameters of detection (confidence and nº of unique peptides) are also indicated

**Table S2. Selected Bio-TT1$_{ct}$-interacting proteins showing significantly increased levels after NMDA treatment compared to BDNF stimulation**

| Uniprot ID | Gene name | logFC NMDA vs BDNF | p-value NMDA vs BDNF | Description | Confidende | nº of unique peptides |
|---|---|---|---|---|---|---|
| A0A0G2JYD8 | *Capn5* | 1.827 | 0.008 | Calpain 5 OS=Rattus norvegicus OX=10116 GN=Capn5 PE=1 SV=2 | High | 3 |
| A0A8I5ZQN4 | *Myo1c* | 0.906 | 0.035 | Myosin 1C OS=Rattus norvegicus OX=10116 GN=Myo1c PE=1 SV=1 | High | 7 |
| A0A8I6G6A0 | *Gna11* | 1.596 | 0.039 | G protein subunit alpha 11 OS=Rattus norvegicus OX=10116 GN=Gna11 PE=1 SV=1 | High | 4 |
| A0A8L2R9X1 | *Abhd6* | 2.091 | 0.024 | Abhydrolase domain containing 6, acylglycerol lipase OS=Rattus norvegicus OX=10116 GN=Abhd6 PE=1 SV=1 | High | 2 |
| A2VCW8 | *Septin7* | 0.863 | 0.042 | Septin OS=Rattus norvegicus OX=10116 GN=Septin7 PE=1 SV=1 | High | 10 |
| B2GUZ3 | *Mthfd1l* | 1.816 | 0.045 | formate--tetrahydrofolate ligase OS=Rattus norvegicus OX=10116 GN=Mthfd1l PE=1 SV=1 | High | 5 |
| D4A280 | *Pak5* | 2.383 | 0.046 | Serine/threonine-protein kinase PAK 5 OS=Rattus norvegicus OX=10116 GN=Pak5 PE=1 SV=1 | High | 2 |

Uniprot ID and gene name are presented for each protein, together with its logFC obtained after comparison of NMDA vs BDNF conditions. Individual p-values for each protein are also shown. Protein description together with quality parameters of detection (confidence and nº of unique peptides detected) are also indicated

**Table S3. Selected Bio-TT1$_{ct}$-interacting proteins showing significantly altered levels after NMDA treatment compared to the basal conditions**

| Uniprot ID | Gene name | logFC NMDA vs Basal | p-value NMDA vs Basal | Description | Confidende | nº of unique peptides |
|---|---|---|---|---|---|---|
| A0A1W2Q6D9 | *Otud6b* | 1.158 | 0.027 | ubiquitinyl hydrolase 1 OS=Rattus norvegicus OX=10116 GN=Otud6b PE=1 SV=2 | High | 2 |
| A0A8I5ZN55 | *Ago2* | -1.303 | 0.048 | Protein argonaute-2 OS=Rattus norvegicus OX=10116 GN=Ago2 PE=1 SV=1 | High | 6 |
| A0A8I5ZNV8 | *Pcnp* | -1.018 | 0.034 | PEST proteolytic signal-containing nuclear protein OS=Rattus norvegicus OX=10116 GN=Pcnp PE=1 SV=1 | High | 4 |
| A0A8I6A326 | *Rbm6* | -1.589 | 0.034 | RNA binding motif protein 6 OS=Rattus norvegicus OX=10116 GN=Rbm6 PE=1 SV=1 | High | 3 |
| A0A8J8YFZ4 | *Ptbp1* | -2.049 | 0.013 | Polypyrimidine tract-binding protein 1 OS=Rattus norvegicus OX=10116 GN=Ptbp1 PE=1 SV=1 | High | 6 |
| A0A8L2R9X1 | *Abhd6* | 1.798 | 0.033 | Abhydrolase domain containing 6, acylglycerol lipase OS=Rattus norvegicus OX=10116 GN=Abhd6 PE=1 SV=1 | High | 2 |
| D4A280 | *Pak5* | 2.460 | 0.038 | Serine/threonine-protein kinase PAK 5 OS=Rattus norvegicus OX=10116 GN=Pak5 PE=1 SV=1 | High | 2 |

Uniprot ID and gene name are presented for each protein, together with its logFC obtained after comparison of NMDA vs basal conditions. Individual p-values for each protein are also shown. Protein description together with quality parameters of detection (confidence and nº of unique peptides detected) are also indicated

**Table S4.** ***Arhgdia*** **(RhoGDI1) and the group of selected Bio-TT1$_{ct}$-interacting proteins included in the Reactome function "Signaling by RhoGTPases"**

| Uniprot ID | Gene name | Description | Confidende | nº of unique peptides |
|---|---|---|---|---|
| A0A0G2K860 | *Arhgef12* | Rho guanine nucleotide exchange factor 12 OS=Rattus norvegicus OX=10116 GN=Arhgef12 PE=1 SV=2 | High | 3 |
| A0A8I5Y1I7 | *Scfd1* | Sec1 family domain containing 1 OS=Rattus norvegicus OX=10116 GN=Scfd1 PE=1 SV=1 | High | 3 |
| A0A8I5YC36 | *Ranbp2* | RAN binding protein 2 OS=Rattus norvegicus OX=10116 GN=Ranbp2 PE=1 SV=1 | High | 4 |
| A0A8I6AHE2 | *Dync1i1* | Dynein cytoplasmic 1 intermediate chain 1 OS=Rattus norvegicus OX=10116 GN=Dync1i1 PE=3 SV=1 | High | 5 |
| A0A8I6ALK1 | *Nup98* | Nuclear pore complex protein Nup98-Nup96 OS=Rattus norvegicus OX=10116 GN=Nup98 PE=1 SV=1 | High | 2 |
| A0A8I6ALV8 | *Tuba1a* | Tubulin alpha chain OS=Rattus norvegicus OX=10116 GN=Tuba1b PE=3 SV=1 | High | 5 |
| A0A8I6G6K9 | *Elmo2* | Engulfment and cell motility 2 OS=Rattus norvegicus OX=10116 GN=Elmo2 PE=1 SV=1 | High | 2 |
| A0A8I6GBC3 | *Actr2* | Actin related protein 2 OS=Rattus norvegicus OX=10116 GN=Actr2 PE=1 SV=1 | High | 7 |
| A0A8I6GLR6 | *Prex1* | Phosphatidylinositol-3,4,5-trisphosphate-dependent Rac exchange factor 1 OS=Rattus norvegicus OX=10116 GN=Prex1 PE=4 SV=1 | High | 2 |
| A0A8L2QIZ5 | *C1qbp* | Complement component 1 Q subcomponent-binding protein, mitochondrial OS=Rattus norvegicus OX=10116 GN=C1qbp PE=1 SV=1 | High | 5 |
| A0A8L2R7U3 | *Tubb2b* | Tubulin beta chain OS=Rattus norvegicus OX=10116 GN=Tubb2b PE=3 SV=1 | High | 2 |
| B2GV73 | *Arpc3* | Actin-related protein 2/3 complex subunit 3 OS=Rattus norvegicus OX=10116 GN=Arpc3 PE=1 SV=1 | High | 4 |
| D3ZRB3 | *Rhof* | RCG21806 OS=Rattus norvegicus OX=10116 GN=Rhof PE=1 SV=2 | High | 2 |
| D4A280 | *Pak5* | Serine/threonine-protein kinase PAK 5 OS=Rattus norvegicus OX=10116 GN=Pak5 PE=1 SV=1 | High | 2 |
| G3V6S0 | *Sptbn1* | Spectrin beta chain OS=Rattus norvegicus OX=10116 GN=Sptbn1 PE=1 SV=4 | High | 91 |
| P68035 | *Actc1* | Actin, alpha cardiac muscle 1 OS=Rattus norvegicus OX=10116 GN=Actc1 PE=2 SV=1 | High | 3 |
| Q5XI73 | *Arhgdia* | Rho GDP-dissociation inhibitor 1 OS=Rattus norvegicus OX=10116 GN=Arhgdia PE=1 SV=1 | High | 8 |
| Q7TT49 | *Cdc42bpb* | Serine/threonine-protein kinase MRCK beta OS=Rattus norvegicus OX=10116 GN=Cdc42bpb PE=1 SV=1 | High | 24 |

Uniprot ID and gene name are presented for each protein together with its description and quality parameters of detection (confidence and nº of unique peptides detected)

### KEY RESOURCES TABLE

| REAGENT or RESOURCE | SOURCE | IDENTIFIER |
|---|---|---|
| Antibodies | | |
| Rabbit polyclonal anti-**C3d** | Dako | Cat#A0063<br>RRID: AB_578478 |
| Mouse monoclonal anti-**CD68 (ED1)** | Millipore | Cat#MAB1435;<br>RRID: AB_177576 |
| Rabbit polyclonal anti-**pS133-CREB** | Millipore | Cat#06-519;<br>RRID:AB_310153 |
| Rabbit monoclonal ani-**GFAP** | Millipore | Cat#MAB360<br>RRID:AB_11212597 |
| Mouse monoclonal anti-**GFAP** | Millipore | Cat#G6171<br>RRID:AB_1840893 |
| Rabbit polyclonal anti-**Iba1** | Wako | Cat#019-19741<br>RRID: AB_839504 |
| Mouse monoclonal anti-**MEF2D** | BD Biosciences | Cat# 610774,<br>RRID:AB_398095 |
| Mouse monoclonal anti-RBFOX3/**NeuN** | Novus | Cat#NBP1-92693SS;<br>RRID:AB_1103747 |
| Rabbit polyclonal anti-neuronal-specific enolase (**NSE**) | Millipore | Cat#AB951;<br>RRID:AB_92390 |
| Rabbit polyclonal anti N-ter region of **RhoGDI** | Santa Cruz | Cat#sc-360<br>RRID: AB_2227516 |
| Rabbit polyclonal anti-**TrkB-FL (C-ter region)** | Santa Cruz | Cat#sc-11;<br>RRID:AB_632554 |
| Rabbit polyclonal anti-**TrkB-T1** (isoform-specific C-ter) | Custom made | |
| Goat anti-mouse IgG Alexa Fluor 546 | Molecular Probes | Cat#A-11030;<br>RRID:AB_144695 |
| Goat anti-mouse IgG Alexa Fluor 647 | Molecular Probes | Cat#A-21236<br>RRID:AB_2535805 |
| Goat anti-rabbit IgG Alexa Fluor 488 | Molecular Probes | Cat# A11034<br>RRID: AB_2576217 |
| Goat anti-rabbit IgG Alexa Fluor 546 | Molecular Probes | Cat# A11035<br>RRID: AB_143051 |
| Donkey anti-rabbit IgG-heavy and light chain-HRP | Bethyl | Cat#A120-108P;<br>RRID:AB_10892625 |
| Donkey anti-mouse IgG-heavy and light chain-HRP | Bethyl | Cat#A90-137P;<br>RRID:AB_1211460 |
| Chemicals and Recombinant BDNF | | |
| **Ara C** (used at 10 µM) | Sigma-Aldrich | Cat#C1768;<br>CAS: 147-94-4 |
| Recombinant Human/Murine/Rat **BDNF**<br>(used at 100 ng/ml) | PeproTech | Cat#450-02 |
| **DAPI** (used at 0.5 or 5 µg/ml) | Molecular Probes | Cat#D1306 |
| **Fluorescein Avidin** D | Vector Laboratories | Cat#A 2001 |
| **Glycine** (used at 10 µM) | Bio-Rad | Cat#161-0718 |
| **Laminin** (used at 4 µg/ml) | Sigma-Aldrich | Cat#L2020;<br>CAS: 114956-81-9 |
| **MTT** (used at 0.5 mg/ml) | Sigma-Aldrich | Cat#M5655;<br>CAS: 298-93-1 |
| **NMDA** (used at 100 µM) | Tocris | Cat#0114;<br>CAS: 6384-92-5 |
| **PhosSTOP** phosphatases inhibitor cocktail tablets | Roche | Cat#04 906 837 001 |
| **Poly-L-Lysine** (used at 100 µg/ml) | Sigma-Aldrich | Cat#P1524;<br>CAS: 25988-63-0 |

| | | |
|---|---|---|
| **Prolong** Diamond antifade reagent | Molecular Probes | Cat#P36970 |
| Complete **protease inhibitor** cocktail tablets | Roche | Cat#11 697 498 001 |
| **Rose Bengal** (used at 20 mg/kg) | Sigma-Aldrich | Cat#R3877;<br>CAS: 632-69-9 |
| **Streptavidin** resin | GenScript | Cat#L00353 |
| **TTC** (used at 2%) | Sigma-Aldrich | Cat#T8877;<br>CAS: 298-96-4 |
| Peptides | | |
| **Bio-sTT1**$_{ct}$ (Biotin-*YGRKKRRQRRR*FVLFHKIPLDG) | GenScript | N/A |
| **Bio-TMyc** (Biotin-*YGRKKRRQRRR*AEEQKLISEEDLLR) | GenScript | N/A |
| **Bio-TT1**$_{ct}$ (Biotin-*YGRKKRRQRRR*PPFVLFHKIPLDG) | GenScript | N/A |
| **TMyc** (*YGRKKRRQRRR*AEEQKLISEEDLLR) | GenScript | N/A |
| **TT1**$_{ct}$ (*YGRKKRRQRRR*PPFVLFHKIPLDG) | GenScript | N/A |
| Critical Commercial Assays | | |
| BCA Protein Assay Kit | Thermo Fisher | Cat# 23225 |
| Clarity Western ECL Blotting Substrate | BioRad | Cat# 1705060 |
| Lipofectamine 2000 | Life Technologies | Cat#11668019 |
| Experimental Models: Organisms/Strains | | |
| Balb/c inbred mice (Balb/cOlaHsd) | Harlan Laboratories | N/A |
| Wistar Rat embryos (E18) | In site facility | N/A |
| Recombinant DNA | | |
| **pCRE** (25-mer oligonucleotide with the sequence of two TrkB CREs subcloned into pTK-Luc) | (Deogracias et al.,2004) | N/A |
| **pMEF2** (pRSRF; -307 to -242 of *Nur77* promoter with two MEF2 sites subcloned into pGL2-basic) | (Woronicz et al., 1995) | N/A |
| **pMEF2mut** (two inactivating point mutations in each MEF2 site of pMEF2) | (Woronicz et al., 1995) | N/A |
| Software and Algorithms | | |
| Adobe InDesign | | RRID:SCR_021799 |
| Adobe Photoshop | | RRID:SCR_014199 |
| GraphPad Prism | | RRID:SCR_002798 |
| ImageJ | https://imagej.net/ | RRID:SCR_003070 |
| R Project for Statistical Computing | http://www.r-project.org | RRID:SCR_001905 |
| String | http://string.embl.de/ | RRID:SCR_005223 |
| Other | | |
| Tissue-Tek O.C.T Compound | Sakura | Cat#4583 |
| Protran Western blotting nitrocellulose membrane | GE Healthcare | Cat#GE10600002;<br>CAS: 9004-70-0 |
